# Modeling cellular self-organization in strain-stiffening hydrogels

**DOI:** 10.1101/2023.12.21.572812

**Authors:** A.H. Erhardt, D. Peschka, C. Dazzi, L. Schmeller, A. Petersen, S. Checa, A. Münch, B. Wagner

**Affiliations:** Weierstrass Institute, Mohrenstr. 39, Berlin, 10117, Germany; Julius Wolff Institute, Berlin Institute of Health, Charité Universitätsmedizin, Augustenburger Platz 1, Berlin, 13353, Germany; Berlin-Brandenburg School for Regenerative Therapies, Augustenburger Platz 1, Berlin, 13353, Germany; IH Center for Regenerative Therapies, Berlin Institute of Health, Charité Universitätsmedizin, Augustenburger Platz 1, Berlin, 13353, Germany; Mathematical Institute, University of Oxford, Woodstock Road, Oxford, OX2 6GG, State, UK

**Keywords:** Hydrogels, nonlinear elasticity, phase-field models, agent-based modeling, cell migration

## Abstract

We develop a three-dimensional mathematical model framework for the collective evolution of cell populations by an agent-based model (ABM) that mechanically interacts with the surrounding extra-cellular matrix (ECM) modeled as a hydrogel. We derive effective two-dimensional models for the geometrical set-up of a thin hydrogel sheet to study cell-cell and cell-hydrogel mechanical interactions for a range of external conditions and intrinsic material properties. We show that without any stretching of the hydrogel sheets, cells show the well-known tendency to form long chains with varying orientations. Our results further show that external stretching of the sheet produces the expected nonlinear strain-softening or stiffening response, with, however, little qualitative variation of the overall cell dynamics for all the materials considered. The behavior is remarkably different when solvent is entering or leaving from strain softening or stiffening hydrogels, respectively.

## 1 Introduction

Mechanical cell-cell interactions as well as mechanical interactions between individual cells and their surrounding extracellular matrix (ECM) play an important role in many biological processes such as tissue regeneration, angiogenesis, cancer and tissue morphogenesis in general [1–6]. Similar to cell migration due to chemotaxis, where cells migrate toward the direction of increasing concentrations of a chemo-attractant, in durotaxis cells migrate along gradients of the ECM stiffness. Typically, cells move towards a higher stiffness [7], while migration towards softer regions has also been observed *in vitro* [8]. Mechanical properties such as viscoelasticity, and nonlinear elasticity directly affect cell migration, where cells respond by restructuring parts of their cytoskeleton [9, 10]. On the other hand, cells actively influence ECM properties like stiffness, or actively remodel the ECM itself [11, 12], which in turn affects cell functions. This interplay between cells and the ECM is fundamental for understanding tissue formation and its properties. Thus, investigating how the temporal mechanical interactions between cells and the ECM influence cellular organization is a very active field of research. As emphasized in [13], the development of hydrogel-based biomaterials is a promising approach for the discovery of new strategies for tissue engineering and regenerative medicine. Thus, it is important to understand how cells mechanically interact with hydrogels and the dependency of cell activity on the properties of the hydrogel. [13] also motivates applying external strains to explore the mechanosensing of cells.

One mode of cell migration is initiated by the adhesion of a cell to the network fibers of the ECM, followed by the cell’s polarization due to mechanical strains of the ECM accompanied by the cytoskeleton restructuring and elongation to initiate cell migration. The internally generated traction force applied by the cells result into a migration path that is either stochastic or follows directional migration paths, depending on various mechanical properties of the ECM [5, 14]. Moreover, while the strain they impose is strongest close to the cell and introduces locally a stiffness that can be many times higher than the applied stress, they also impact their surrounding region [15]. As shown in Liu et al. [16], cells, or even a single cell, may induce bands of fibers that lead to an increase in fiber density and alignment between cells. Such mechanical coupling between cells is suggested to be a universal mechanism in natural ECM and in particular collagen and fibrin gels [17]. The strain-stiffening property is characteristic of the ECM and typical for natural hydrogels, i.e. composed of biomolecular networks, such as actin, type I collagen [18], fibrin [17]. It has been measured by Han et al. [19], where microrheology was used to measure the nonlinear stiffening of fibrin matrices induced by single cells. To some extent nonlinear stiffening has even been observed experimentally for stiff tissues such as cortical bones [20].

Apart from the details of the impact of microstructure of the ECM on cell-ECM mechanical interactions, which is reviewed in [3, 21, 22], we will focus here on strain-stiffening properties by introducing a Gent-type model into our hydrogel model for the ECM.

Many ECM materials can be experimentally characterized and mathematically modelled as viscoelastic hydrogels [23]. In the past, hydrogels have been used in cell culture to study various cell functions, and to isolate mechanisms influencing cell migration [24]. Also, for comparison, stress-strain measurements can in principle be directly performed under stress- or strain-controlled conditions to distinguish between the elastic and viscous components. In addition, stress change over time under a specific strain allows to infer details of hydrogel relaxation, which can give insights if applied traction forces lead to force or dissipation of the energy as a response [25–27]. Insight gained from such comparisons could potentially improve our understanding of specific cell-ECM interactions and help design new hydrogel materials and develop strategies to control cell migration.

One main goal of our theoretical study is to investigate how initial seedings of cells on the ECM evolve into self-organized cell patterns by taking into account the local and long-range mechanical interactions and solvent diffusion. To this end, we use agent-based modeling, a versatile framework in which discrete cells can be simulated to behave autonomously according to a set of rules that can represent events at different levels [28]. Agent-based cellular models coupled with finite element model describing the ECM are commonly utilized to model and investigate phenomena like bone regeneration, see [29, 30] and the references contained therein. In this line of research, we study cell-hydrogel and cell-cell interactions for stretched and unstretched hydrogels, cf. again [13], and we investigate the long-time evolution of an interacting cell system. The impact of mechanical ECM properties is investigated by comparing strain-softening neo-Hookean and strain-stiffening Gent-type behavior and we allow for solvent exchange with the surrounding. Performing a plane-strain or a plane-stress approximation, we study two effective 2D models for thin sheets or thick elastic layers.

The paper is structured as follows: In Sec. 2 we derive the mathematical model framework for the cells and the hydrogel, beginning with the derivation of a thermodynamically consistent continuum model for the hydrogel as a two-phase system consisting of a nonlinearly elastic network in a liquid solvent in 3D in Sec. 2.1. For later comparison we employ two different nonlinear elasticity models, a neo-Hooke and a Gent-type model, the latter accounting for strain-stiffening properties of the fiber network. This is followed by the derivation of an approximation for a thin hydrogel sheet and specification of the fixed and free boundary boundary conditions. We then introduce the agent-based model (ABM) in Sec. 2.3, specify the migration rules and the iterative path of migration decisions by the cells when interacting with the hydrogel sheet.

In Sec. 3 we formulate the problem for a thin hydrogel sheet and investigate scenarios of a sheet for a range of applied loads or stresses. Since the dynamic and morphological evolution of stretched hydrogel sheets has not been modeled before, at least not as a continuum multi-phase free boundary system, this analysis is necessary to understand the response in form of stress-strain relations and the corresponding temporal and spatial evolution for different parameter configurations, different elasticity models, such as neo-Hooke or Gent, and different solvent concentrations. Here, we also investigate the effects of possible solvent loss (or gain, i.e. swelling). This will serve as a guidance for investigating the collective response of the cell population on hydrogel sheets under various scenarios in the following section. For comparison, we also contrast this in this section with corresponding results for the purely elastic counterparts for neo-Hooke and Gent models, since there is a considerable literature using such models, see e.g. [31–33].

Finally, in Sec. 4 we explore the migration response of cell populations for various mechanical stimuli provided by the hydrogel sheet. Binning with relaxed conditions, i.e. no stretching, as a reference state, for hydrogel as well as purely elastic sheets, we vary the magnitude of the applied stretch, flux conditions at the free boundaries, strength of traction forces exerted by the cells. Following the evolution of the cell population we monitor corresponding pattern formation of the cells together with their orientation, distribution of solvent within the hydrogel sheet and the stress state. In particular, we explore conditions for chain formation of cells, that are well documented in the literature and discuss underlying mechanisms for their formation as well effects of strain stiffening. In Sec. 5 we conclude and summarize this work, discuss future applications as well as extensions of our model framework.

## 2 Model framework for cell-hydrogel interaction

We develop a versatile and extensible mathematical framework for viscoelastic hydrogels to describe cell migration in an extracellular matrix. First, we derive a time-dependent continuum model for a hydrogel based on the thermodynamics of the two-phase system consisting of cross-linked biopolymers and a solvent. Then, we perform a model reduction to effective 2D hydrogels models in order to consider thin hydrogel sheets. Finally, we develop the corresponding ABM for cell motion on thin sheets.

### 2.1 Hydrogel model

First, we formulate a minimal hydrogel model that exhibits some typical mechanical properties of an ECM using a class of hydrogels that consists of a cross-linked network of biomolecules, solved in an aqueous solution. For this we derive a time-dependent two-phase continuum model of coupled partial differential equations in 3D describing the evolution of two species (solvent and biomolecular network), where the mechanical properties are encoded in the mechanical energy for the cross-linked network. Our model also accounts for possible solvent exchange with the environment to allow for contraction or swelling via free boundaries, reflecting the typical aqueous environment of an ECM. In fact, other material properties such as growth factors can be modeled as well and are introduced using additional phases or components.

The elastic behavior of most hydrogels of synthetic polymers such as polyethylene glycol (PEG) and polyacrylamide (PA) is similar to that of rubber-like materials. Their elasticity can be understood on the basis of the classic theory of rubber elasticity. For biopolymer hydrogels such as actin, collagen and fibrin, rheological properties such as strain-stiffening have to be incorporated in the hydrogel model. This entails that at small deformations biopolymer hydrogels display a constant stiffness or storage modulus that depends on polymer concentration. When the applied stress (external or internal) is large enough they show a strong nonlinear response, with much higher values than the original elastic modulus.

The dynamic hydrogel model is formulated in the Lagrangian frame Ω ⊂ ℝ^3^. The mechanical state of the hydrogel is characterized by a function ***χ*** : [0, *T*] × Ω→ ℝ^3^, where ***χ***(*t*, ***x***) ∈ ℝ^3^ encodes the deformed position of a material point ***x*** ∈ Ω at the time *t* ∈ [0, *T*]. With ***F*** = ∇***χ*** ∈ℝ^3*×*3^ we denote the deformation gradient and with *J* = det(***F***) its determinant. The chemical state of the material is characterized by a concentration ***c*** : [0, *T*] × Ω → ℝ^*n*^, where each component of ***c*** = (*c*_1_, …, *c*_*n*_) encodes the concentration *c*_*i*_ of a component *i* = 1,…, *n* for *n* ∈ ℕ. Combining deformation and concentration into a state vector ***q*** = (***χ, c***) we have a free energy functional for the hydrogel

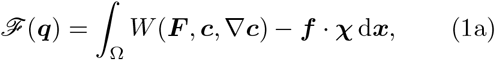

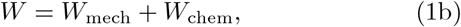

where the density *W*_mech_(***c, F***) contains the elastic stored energy of the hydrogel network and the density *W*_chem_(***F***, ***c*, ∇*c***) contains chemical and entropic contributions to the free energy that drive diffusion, phase transitions and phase separation and are therefore thermodynamic in nature. Additionally, with ***f*** : Ω →ℝ^3^ we encode given (external) mechanical forces. Each component takes up a certain volume fraction *φ*_*i*_ = *α*_*i*_*c*_*i*_*/J* with the component *α*_*i*_ *>* 0 of the vector of specific molar volumes *α* ∈ ℝ^*n*^. Setting

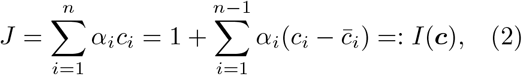

we have by construction 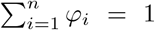. In the last step we assumed that *c*_*n*_ corresponding to the polymer network concentration is constant in space and time and that *J* = 1 in the reference state, where 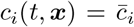 for *i* = 1, …, *n* − 1 and ***x*** ∈ Ω. This constraint is enforced by a Lagrange multiplier *p* : [0, *T*] × Ω → ℝ contained in a state vector ***q***_*p*_ = (***q*** = (***χ, c***), *p*) using the Lagrangian

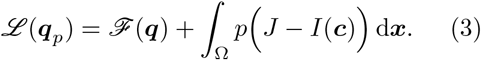

Minimizers of the free energy (1) subject to the incompressibility constraint (2) can be obtained by considering saddle points of the Lagrangian (3). Now, we construct a thermodynamic consistent evolution law for this system, i.e. the free energy is decreasing over time. Thus, we introduce chemical potential-like variables 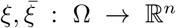 and construct the diffusive mobility of components

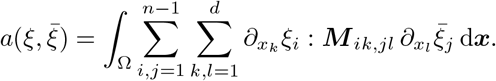

In general, ***M*** (***q***) is a non-negative tensor that acts on components and spatial indices and depends on the concentration ***c***. A further dependence of ***M*** on ***F*** allows to include both anisotropic diffusion and Eulerian interfacial energies. The corresponding diffusion is often diagonal in the concentration and with mobility function *m*_*i*_ = *m*_*i*_(*c*_*i*_) leads to

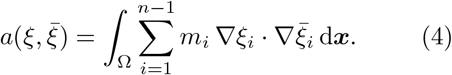

Corresponding to the time derivative of the state *∂*_*t*_***q***_*p*_ = (*∂*_*t*_***χ***, *∂*_*t*_***c***, *∂*_*t*_*p*) we consider general rates ***w*** = (***w***_*χ*_, *w*_***c***_, *w*_*p*_) with ***w***_*χ*_ : Ω → ℝ^3^, *w*_***c***_ : Ω → ℝ^*n*^ and *w*_*p*_ : Ω → ℝ and define a bilinear form

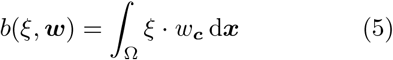

coupling chemical potentials *η* : [0, *T*] × Ω→ ℝ^*n*^ and these rates. We model the evolution of the hydrogel by a nonlinear saddle point problem, where we seek ***q***_*p*_(*t*) and *η*(*t*) such that

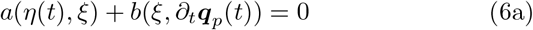

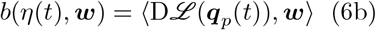

holds for all (test functions) ***w*** and *ξ* at time *t* ∈ (0, *T*). Here, we introduced the (Gateaux) derivative of the Lagrangian functional

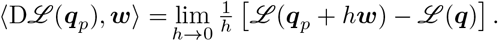

For details of the corresponding model derivation we refer to [34, 35].

Thermodynamic consistency holds for (6) in the sense that if we choose ***w*** = (*∂*_*t*_***χ***, *∂*_*t*_***c***, 0) we get from (6b) that 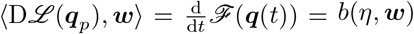, where we used that the constraint (2) is satisfied. Testing the first equation (6a) with the chemical potential *ξ* = *η* we obtain the final result

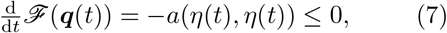

where we used that *a*(*ξ, ξ*) ≥ 0 holds by construction of *a* for any *ξ*, since we consider nonnegative ***M*** in (4). We supplement this problem with homogeneous natural boundary conditions for ***c*** on the entire boundary *∂*Ω. For the deformation we impose homogeneous natural boundary conditions on Γ_*N*_ ⊂ *∂*Ω and inhomogeneous Dirichlet boundary conditions ***χ***(*t*, ***x***) = ***χ***_Γ_(*t*, ***x***) for ***x*** ∈ Γ_*D*_ = *∂*Ω \ Γ_*N*_ and given ***χ***_Γ_. Note that (7) does not strictly hold for time-dependent inhomogeneous Dirichlet boundary conditions, since the elastic energy effectively depends explicitly on time. However, while the free energy might even increase, the model (6) is thermodynamic consistent, since the resulting first Piola–Kirchhoff stress tensor **P** and the chemical potentials *η*

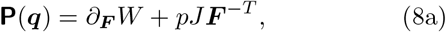

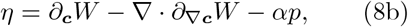

contain all fundamental cross-coupling effects. Note that we used *∂*_***F***_ *J* = *J* ***F*** ^−*T*^. The corresponding Cauchy stress is obtained as ***σ*** = *J* ^−1^**P *F*** ^*T*^.

The resulting general hydrogel evolution is

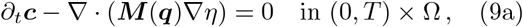

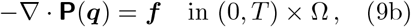

with the constraint (2) and boundary conditions

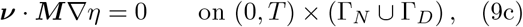

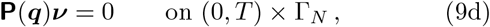

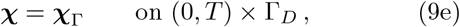

and initial conditions 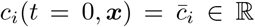 for *i* = 1, …, *n*− 1 and ***χ***_Γ_(*t* = 0, ***x***) = ***x***. Here ***ν*** is the outer normal vector field on *∂*Ω. Alternatively to (9c), we also consider cases where solvent can flow into the gel through the front faces with a flux ***ν***· ***M*** (***q***) ∇*η* + *n*(***q***)*η* = 0, which in an effective 2D model for a thin sheet will lead to volumetric source terms in the hydrogel model. We assume that the system is initially in equilibrium, i.e. *ℱ* is minimal for 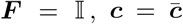and *p* = 0 for ***f*** = 0. Next, we specify the energy density and restrict to two components, i.e. *n* = 2, for solvent and elastic polymer molecules. In order to investigate the impact that the solvent diffusion has on the gel dynamics, we want to be able to revert the model to a simple hyperelastic model. This can be achieved by switching off the diffusion setting ***M*** = 0 and thus 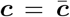 and *J* = 1 for all times. In the following we explain specific choices for the energy density that reflect the expected mechanical and thermodynamic properties of hydrogels.

#### Entropic energy density

The energy density *W*_chem_ in (1) contains chemical and entropic contributions of thermodynamic origin that contribute to the free energy. We use here a Flory-Huggins free energy [36–38] expressed using gradients of the concentration and using volume fractions as

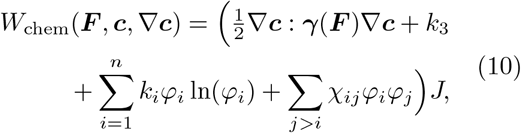

which describes the mixing of the polymeric system [34]. Here, we restrict to a binary mixture with *n* = 2, where 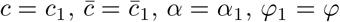 and *φ*_2_ = 1 − *φ* so that the parameters in the density reduce to ***γ***, three constants *k*_1_, *k*_2_, *k*_3_ and the mixing parameter *χ*_12_ such that

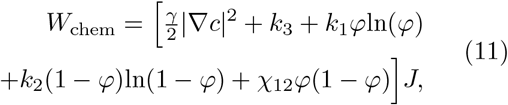

with *φ* = *αc/J*. Depending on the sign of the mixing parameter *χ*_12_ one can trigger whether mixing or phase separation is energetically favorable. In the following we assume *χ*_12_ = 0. While *k*_1_, *k*_2_ *>* 0 can be chosen arbitrarily and 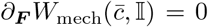 see Figure 1, we set *k*_3_ so that *∂*_***F***_ *W*_chem_ = 0 in equilibrium, i.e. 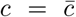 and ***F*** = 𝕀. Therefore, the material is stress-free in the prepared reference state. In [39] a similar constant *k*_3_ was used to ensure existence of minimizers by ensuring *W*_chem_(***F***, ***c***, ∇***c***) ≥ 0 for all det ***F*** ≥ 0.

**Fig. 1.**
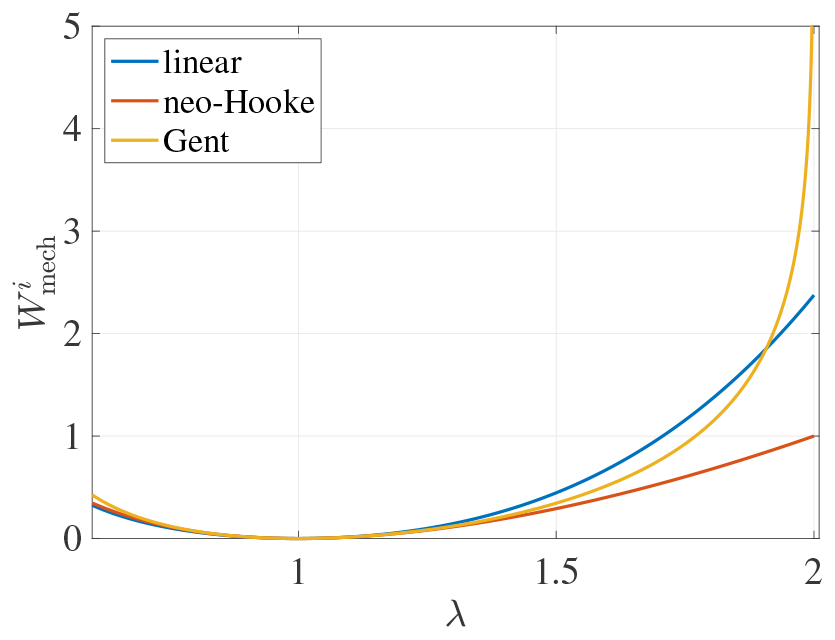
Mechanical energy density with *μ* = 1 for linear elastic material, neo-Hooke and Gent model (*J*_*m*_ = 2) for the deformation in (14). The 3D Gent model becomes singular at *λ* = 2.

#### Elastic energy density

The elasticity of hydrogels can in general be understood on the basis of the classic theory of rubber elasticity such as for synthetic polymers for the network phase. However, the elastic modulus of biopolymer networks in natural hydrogels, such as actin, fibrin or collagen, typically exhibit strain-stiffening properties. For this study we use neo-Hookean elastic materials as one of the simplest nonlinear elasticity models, or alternatively a Gent-type material that accounts for the strain-stiffening properties of a hydrogel. Therefore, in the following we introduce elastic energy densities 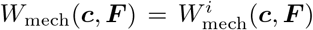 which allows us to investigate properties of strain-softening by a neo-Hookean energy *i* = “neo-Hooke” or strain-stiffening by a Gent-type energy *i* = “Gent”. These nonlinear material models for large deformations contrast a simple linear elastic model typically valid only for small deformations, see Fig. 1.

Commonly used constitutive material laws for describing the nonlinear elastic response of a polymer network are of *neo-Hookean* type [40]. Here, the strain energy density is given by

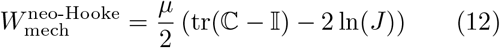

with 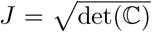 and left Cauchy-Green tensor ℂ = ***F*** ^*T*^ ***F***. Compared to a linear elastic material, the stress-strain relation also starts linear but will flatten for larger strains.

We model strain stiffening hydrogels by employing the *Gent model* [41] with strain energy

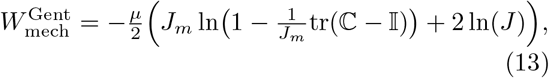

where *J*_*m*_ = *I*_*m*_ − 3 is the so-called limiting deformation of the material and *I*_1_ = tr(ℂ) is the first invariant of the right Cauchy-Green deformation tensor. Stretchability of soft tissue varies with age, pathology, humidity, as well as the type of tissue [42]. Tendons and ligaments can be uniaxially stretched to a strain of around 15% [43], cartilage 120% [43], skins 110% [44] and aorta 100% [43]. These value correspond to a deformation *J*_*m*_ = 0.06, 2.75, 2.36 and 2.00, respectively. This parameter can be tuned by changing the cross-link density of the polymer chains or by varying the mixture of different kinds of polymers. In the limit *I*_*m*_→ ∞ the Gent model converges to the neo-Hookean model. In Fig. 1 we show the three elastic energy densities for a uniaxial deformation with *J* = 1 of the form

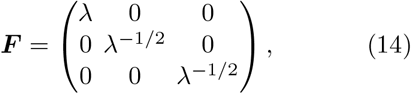

showing clearly that the energies 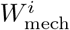 are minimal for ***F*** = 𝕀 and that Gent or neo-Hookean materials are softer or stiffer compared to linear elastic ones for larger strains *λ*, respectively. For elastic models one might want to consider a dependence *μ* = *μ*(***c***) to describe the effect of drying-induced softening or stiffening as in [45], but here we will consider a constant elastic modulus that does not depend on concentration.

### 2.2 Dimension reduction for sheets

We consider an elastic sheet in a rectangular cuboid domain 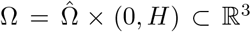 with 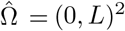 with the simple sheet geometry shown in Fig. 2. In that sketch we introduce front, side, top and bottom faces on which boundary conditions are defined. In order to solve (9) we use inhomogeneous Dirichlet boundary conditions on Γ_*D*_ = Γ_top_ ∪ Γ_bot_ and no-stress boundary conditions on Γ_*N*_ = Γ_side_ ∪ Γ_front_.

**Fig. 2.**
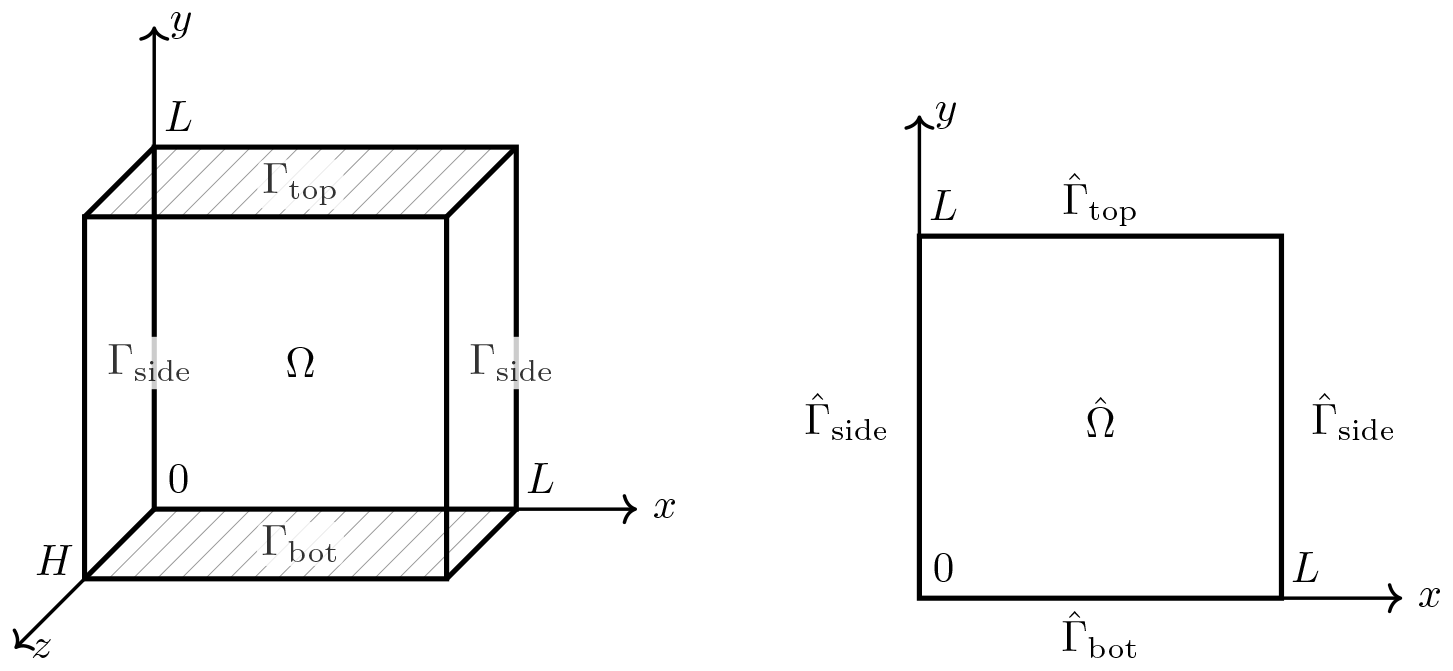
(left) 3D sheet 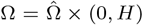) and (right) its 2D cross section 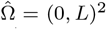. We denote the top and bottom faces of the cuboid by 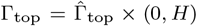 and 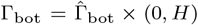 where 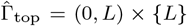 and 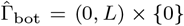, as indicated by the gray dashed lines. The side faces of the cuboid are 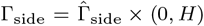 where 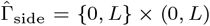. Additionally, the front faces of the cuboid are Γ_front_ = (0, *L*)^2^ × {0, *H*}.

Model reduction for elastic sheets and membranes is usually based on an assumption that a general elastic deformation ***χ***(*t*, ***x***) = ***x*** + ***u***(*t*, ***x***) depending on time *t* and space 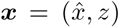with 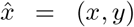 and with displacement ***u***(*t*, ***x***) = (*u*_*x*_(*t*, ***x***), *u*_*y*_(*t*, ***x***), *u*_*z*_(*t*, ***x***)) can be approximated

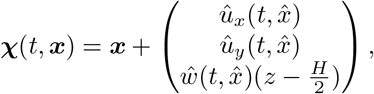

with functions 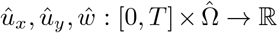 that only depend on the cross-section 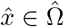 and thus

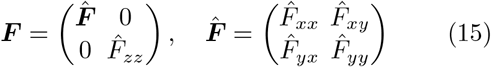

such that 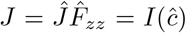 with 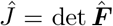 holds for the determinant. Furthermore, we approximated the concentration by 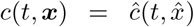. Under these assumptions one distinguishes the plane-stress approximation for thin sheets *H* ≪ *L* and the plane-strain approximation for *H* ≫ *L*.

#### Plane stress approximation

Solving the above condition for a general 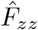 with 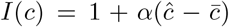 given by the constraint (2) yields an explicit expression

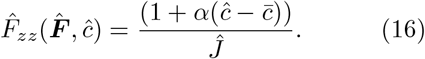

This allows to approximate the original three-dimensional problem by a two-dimensional problem for 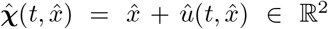 where we use *û* = (*û*_*x*_, *û*_*y*_) and with 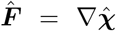 we can replace 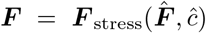 via (15) and (16) in the free energy. This approximation is commonly used for thin sheets *H* ≪ *L* with appropriate boundary conditions, e.g. here we are going to use *u*_*x*_ = *u*_*z*_ = 0 and 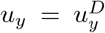 on Γ_*D*_ for the 3D model or correspondingly *û*_*x*_ = 0 and 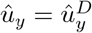 on 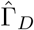 for the corresponding plane stress approximation. Note that in the plane stress approximation the constraint (2) is automatically satisfied and no multiplier is required. Thus, the plane stress approximation of a 3D incompressible material is effectively a 2D model that allows for compression in the *xy*-plane.

#### Plane strain approximation

The corresponding plane strain approximation assumes *ŵ*= 1 and thus

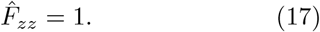

This produces *J* = *Ĵ*, which in the plane stress approximation also needs to be enforced using a Lagrange multiplier. Also this yields 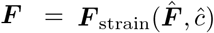 (here trivially independent of *ĉ*) that can be used in the free energy. This approximation is commonly used for *H* ≫ *L* or with appropriate boundary conditions on Γ_front_. Thus, the plane strain approximation of a 3D incompressible material is effectively a 2D model that also does not allow for compression in the *xy*-plane.

#### Models for two-dimensional sheets

In that sense, the main step of both approximations is to replace the energy in (1) by the 2D approximation 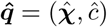 which gives

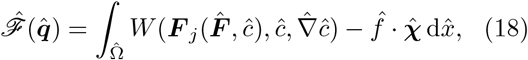

with *W* = *W*_mech_ + *W*_chem_ as before, which relies on the assumption that the external force density was already effectively two-dimensional before, i.e. 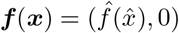. We define 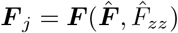 from (15) for *j* = “stress” by inserting 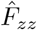 from (16) and for *j* = “strain” by inserting 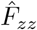 from (17), respectively. While in the plane stress approximation 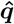 already satisfies the constraint, in the plane strain approximation we need to enforce it via the Lagrange multiplier *p* contained in 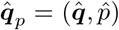 by defining the Lagrange functional

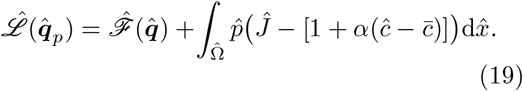

The bilinear forms for the effective 2D model are

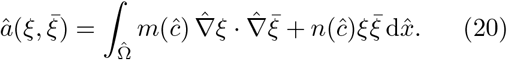

and with rates ***w*** = (*w*_***χ***_, *w*_*c*_) for plane stress or ***w*** = (*w*_***χ***_, *w*_*c*_, *w*_*p*_) for plane strain let

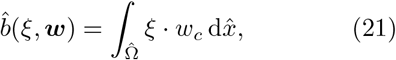

where we avoided the “hats” on chemical potentials and rates for ease of notation. A nonzero mobility *n*(*ĉ*) in (20) leads to a non-conserved evolution caused by a possible contact of the hydrogel with a solvent bath through the side and front faces. Introducing a time discretization 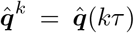 with time step size *τ*, in the plane stress approximation this leads to

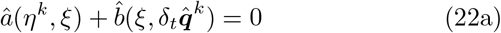

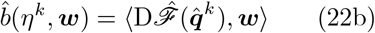

with 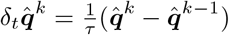 and in the plane strain approximation with

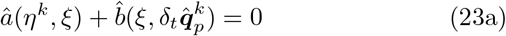

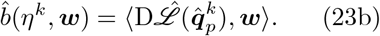

with 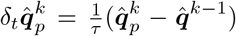. As described in [35], this nonlinear saddle point problem is discretized in space using finite elements on triangular meshes and discretized in time via an incremental minimization. We implement the problem using the finite element library FEniCS [46, 47]. Solving (22) or (23) with a force 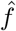 produced by the cell configuration contributes to one time step (iteration) of the hydrogel model. For more details we also refer to the supplementary source code [48]. For thin sheets, a comparison of 2D and 3D deformation and concentration in the plane stress approximation in shown in Fig. 3. While in particular close to the top and bottom boundary, due to a mismatch of boundary conditions in the plane stress approximation slight differences in the solvent concentration are visible. Nevertheless, the 2D deformation describes the thin 3D hydrogel sheet very well. In the following we explain the ABM and the construction of the force 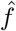 that couples to the hydrogel.

**Fig. 3.**
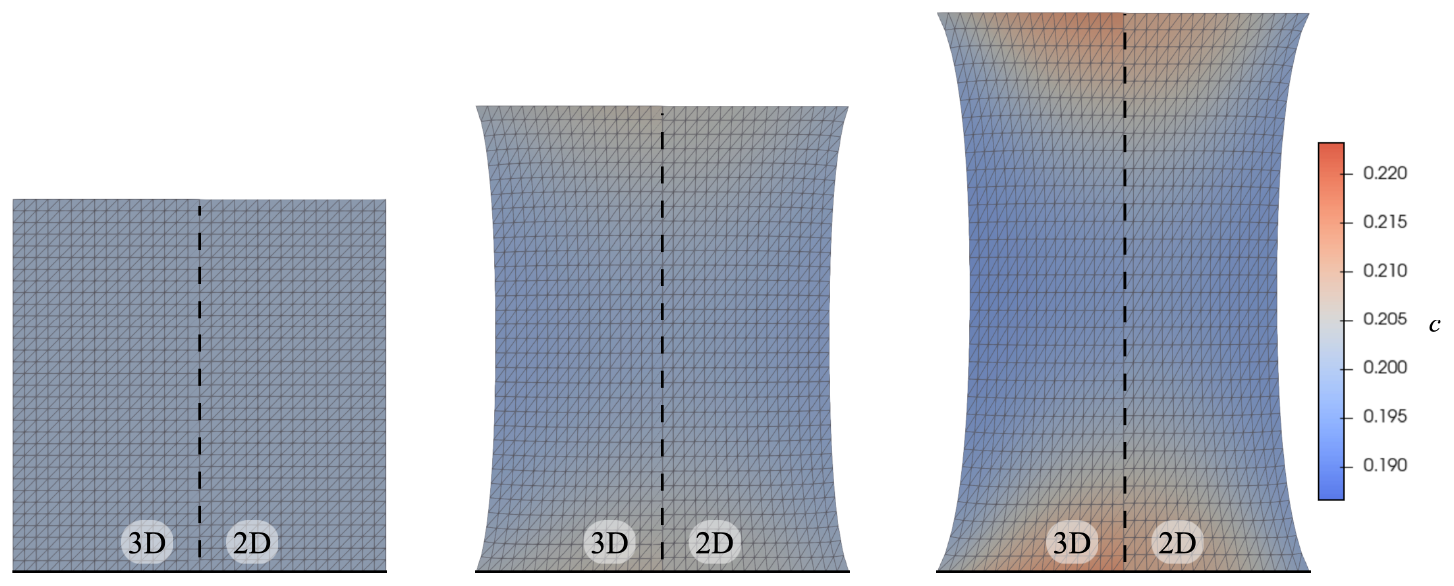
Comparison of 3D hydrogel model with the corresponding 2D hydrogel model in plane stress approximation (Gent). We show the deformed domain ***χ***(Ω) (mesh) and the concentration *ĉ* (shading) for increasing applied strain ***χ***_Γ_ from left to right. The left half of each plot is from the 3D problem and the right half from the effective 2D problem.

### 2.3 ABM for cell migration

For the interaction of the ECM with the cells we neglect the short time adhesion process between cell and hydrogel, and focus on the nonlinear mechanics on the time-scales of cell migration. Cells are characterized by a spatial location and an orientation and each probing the hydrogel by exerting a dipole traction force of a given strength that is transmitted to the hydrogel and in turn to other cells, that are also probing the hydrogel. In our current study we restrict the investigation to a few basic rules, that are generally thought to govern cell migration in cell-ECM systems, namely that cells move to locations of highest stiffness. The collective migration for a given distribution of cells is then described by an ABM governing the migration decisions of each cell as a function of the states of other cells and the hydrogel.

Migrating cells (fibroblasts) are assumed to have an elongated shape as they move through their hydrogel environment by exerting dipole-like traction forces on the hydrogel, pulling two regions to which the cell attaches toward the cell center. Since cells have an elongated shape, each cell also posseses an orientation. While we assume that cells within the thin hydrogel sheet have four possible orientations, i.e. horizontal, vertical or in one of two diagonal orientations, they align and can only be oriented horizontally at the top and bottom boundaries and vertically at the left and right boundaries of the rectangular hydrogel sheet. These properties of cells can be described using the following sets and functions.

#### Cell state space

First, we define the set of admissible states *A* of single cells *a* ∈ *A* and then the set of admissible states ***A*** of many cells 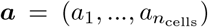 ∈ ***A*** with *a*_*i*_ ∈*A* and *n*_cells_ ∈ℕ. Therefore, for given *N* ∈ℕ and *h* = *L/N* let us construct the sets of cell positions and cell orientations by

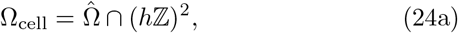

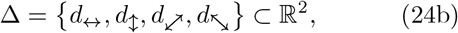

where *d*_↔_ = (*h*, 0), *d*_↕_ = (0, *h*), 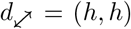 and 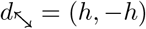. By combining position 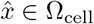 on a lattice and discrete orientations *d* ∈ ∆ in pairs, we build the set of admissible single cell states

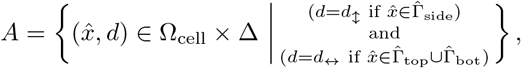

i.e. cells on the boundary are aligned in the sense 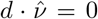 with the normal vector field 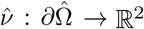. From this we construct the set of multi-cell configurations with *n*_cells_ ∈ ℕ cells by

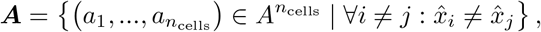

i.e. cells 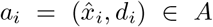 for 1 ≤*i* ≤*n*_cells_ occupy different positions. The cells generate a dipole-type force 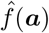 that pulls material towards the cell center by pulling in the direction given by the cell orientation. For any cell configuration 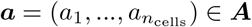 that force is

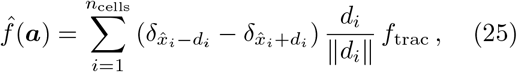

where the traction strength is *f*_trac_ *>* 0, see also Fig. 5. For any given state of the hydrogel 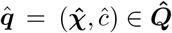, a single cell 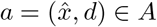 in the ABM can probe the response generated by the traction force by the function 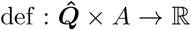 that measures the local deformation at the cell location 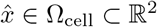 via

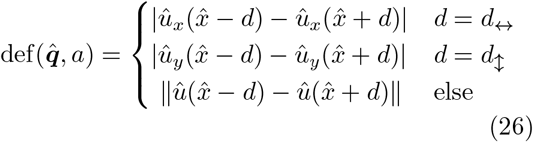

with 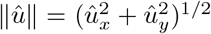 and where *û* = (*û*_*x*_, *û*_*y*_). This function does not explicitly depend on *ĉ*. The state of the ABM for cell motion at any discrete time *t* = *kτ* is given by ***a***^*k*^ ∈ ***A***. The definitions before now allow us to model the dynamic evolution of the cell configuration ***a***^*k*^ and the hydrogel 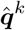 by updating 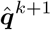 and ***a***^*k*+1^ at each time step.

#### Multi-cell dynamics

The final modeling step is a description of the cell migration via an ABM satisfying certain rules [5, 6, 49, 50]. In the first step, the ABM is *seeding the biological cells* on the substrate, e.g. an elastic material or a hydrogel, cf. Figure 4.

**Fig. 4.**
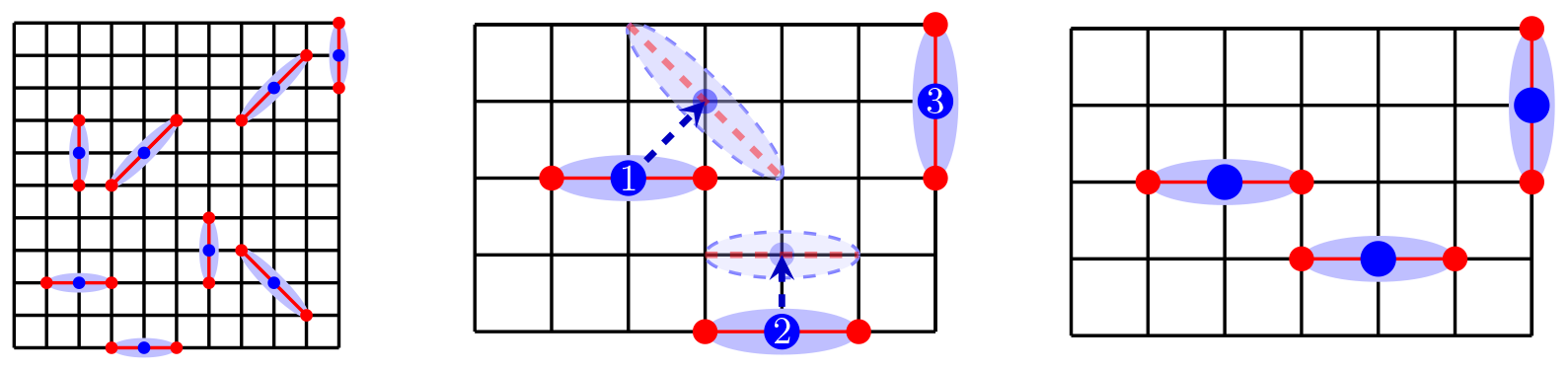
**(left)** Exemplary multi-cell state 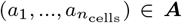 for a lattice of cell position Ω_cell_ with *N* = 10 and *n*_cells_ = 8 and 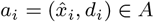. The cell center positions 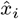 are indicated using blue dots and the corresponding orientations *d*_*i*_ are indicated using red dots at 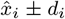 and red lines connecting these with the cell center. For any given state ***a***, the traction force 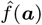 pulls the hydrogel at each red dot towards the corresponding cell center with strength *f*_trac_. **(middle)** Proposed cell update for the cell state from the bottom left corner of the multi-cell state. The new positions and orientations for cell 1 and cell 2 (white labels) are indicated using light blue dots and dashed blue lines, whereas cell 3 is fixed because at every iteration not all cells are updated. **(right)** New configuration where the updated cell state was rejected for cell 1 and accepted for cell 2.

**Fig. 5.**
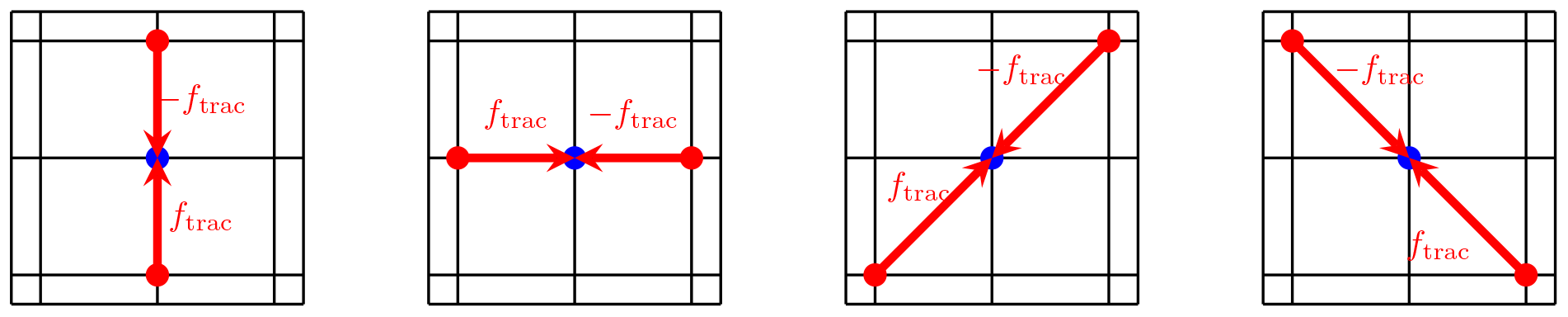
Traction forces with a magnitude *f*_trac_ on the ECM induced by the cell (blue dot).

Cells act on the ECM by exerting a dipole-type traction force. The dipole orientation is given by the cell orientation and pulls material towards the cell core, cf. Fig. 5. Each time step, approximately 80% of the cells are moved to a neighboring cell position. The cells on the left and right boundaries can only apply traction forces as vertical dipoles, while the cells on the top and bottom can only apply traction forces as horizontal dipoles. All other cells can also apply traction forces as diagonal dipoles. The cells have a clear preference to stay on a stiff place [51], i.e. cells prefer a locations with small deformation. Therefore, after moving the cells to a random adjacent cell position and assuming a random orientation, the algorithm determines the deformation of the substrate at each cell based on the mechanical deformation and the dipole orientation. In case the new proposed cell position is less stiff compared to the old cell position, i.e. produces a higher deformation as a reaction to the dipole force, the cell moves back provided the previous cell position is unoccupied. Then, the new deformation of the ECM is determined by solving the hydrogel model with traction forces generated by the new cells configuration with cells that migrated to potentially stiffer positions. We summarize this procedure in Table 1.

**Table 1.**
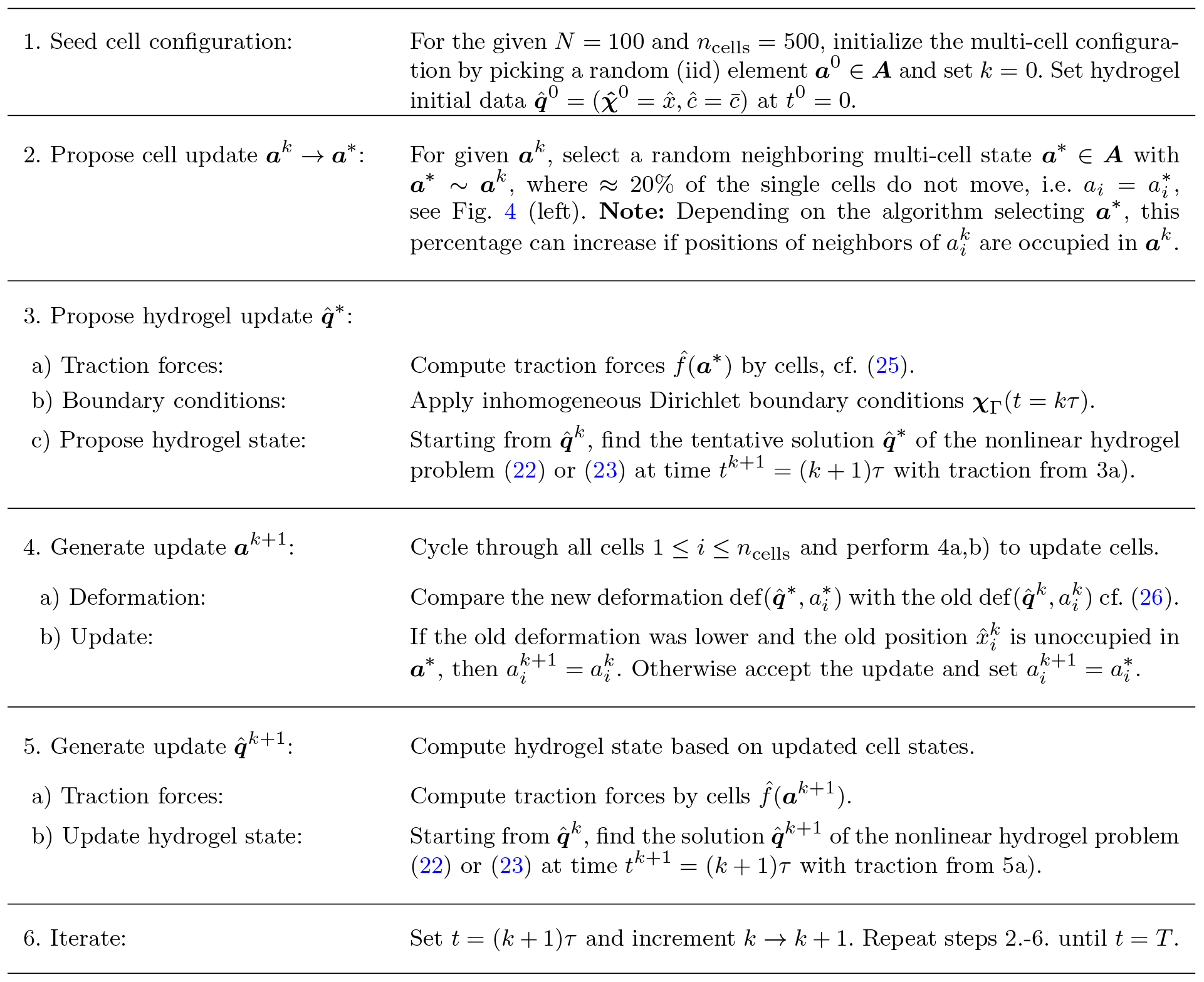
Implementation of ABM that computes ***a***(*t*^*k*+1^) = ***a***^*k*+1^ and 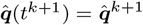 based on the solution at previous time *t*^*k*^ = *kτ*. The update is performed via intermediate states for cells ***a***^∗^ and for hydrogel 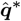. We say two single cells 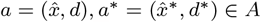 are *neighbors* and write *a* ∼ *a*^∗^ if there exists a direction 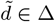 such that 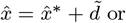 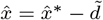 or if *a* = *a*^∗^. Two multi-cell configurations ***a, a***^∗^ ∈ ***A*** are *neighbors* and we write ***a*** ∼ ***a***^∗^, if all of their components are neighbors 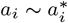 for *i* = 1, …, *n*_cells_. We say a cell position 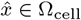 in 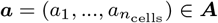 is *unoccupied* (by a cell), if for no component 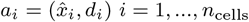 we have 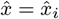. Otherwise we say the position 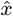 is *occupied* (by a cell) in ***a***.

**Table 2.** Model-specific mathematical notation and explanation of symbols.

| agent-based model |  |  | hydrogel model |  |  |
| --- | --- | --- | --- | --- | --- |
| symbol | name | def. | symbol | name | def. |
| $\hat{x}$ | cell position | Sec. 2.3 | $\varphi$ | volume fraction | Sec. 2.1 |
| $d$ | cell orientation | Sec. 2.3 | $\mathbf{c}$ | concentration | Sec. 2.1 |
| $n_{\text{cells}}$ | number of cells | Sec. 2.3 | $\chi$ | deformation | Sec. 2.1 |
| $A$ | single-cell state | Sec. 2.3 | $\mathbf{F}$ | deformation gradient | Sec. 2.1 |
| $\mathbf{A}$ | multi-cell state | Sec. 2.3 | $J$ | Jacobian determinant | Sec. 2.1 |
| $\Omega_{\text{cell}}$ | cell position set | Eq. (24a) | $\mathbf{P}$ | first Piola-Kirchhoff stress | Eq. (8a) |
| $\Delta$ | cell orientation set | Eq. (24b) | $\mathbf{q}$ | state vector | Eq. (1) |
| def | deformation function | Eq. (26) | $\chi_{\Gamma}$ | Dirichlet boundary cond. | Eq. (27) |
| $\hat{f}$ | traction force | Eq. (25) | $\mathbf{x}$ | coordinates $(\hat{x}, z) \in \mathbb{R}^3$ | Sec. 2.1 |
| $\delta_x$ | Dirac δ-distribution | | $\hat{x}$ | coordinates $(x, y) \in \mathbb{R}^2$ | Sec. 2.2 |
| $f_{\text{trac}}$ | traction strength | Eq. (25) | $\mathcal{F}$ | free energy functional | Eq. (1) |
| $a_i \sim a_j$ | neighboring cells | Table 1 | $\mathcal{L}$ | Lagrangian functional | Eq. (3) |
| $N$ | cell grid size | Eq. (24a) | $W$ | energy density | Sec. 2.1 |
| $N_{\text{dipole}}$ | dipole count | Eq. (31) | $\mathbb{I}_d$ | $d \times d$ identity matrix | |
| | | | $\mathbb{C}$ | left Cauchy-Green tensor | Sec. 2.1 |
| | | | $I(\mathbf{c})$ | incompressibility constraint | Eq. (2) |

## 3 Strain-stiffening effects in stretched hydrogel sheets

Before we investigate the interactions of cells on hydrogel sheets and emerging population pattern, we focus on the material properties of hydrogel sheets for a range of applied stresses. In particular we are interested in the different mechanical response for hydrogels, where the nonlinear elastic energy density is of neo-Hooke or of Gent type. The comparison will give insight into the parameter ranges when to expect significant differences in cell behavior as properties such as strain-stiffening will become relevant. To this end, we present a brief comparative study regarding the material properties under plane strain and plane stress condition discussed in Sec. 2.2, respectively, while stretching their elastic sheets.

We note, that mechanical and morphological properties of elastic sheets is a fascinating research topic by itself and has been investigated in the past, in particular for purely elastic sheets [52, 53], where however the strain-stiffening aspects have not been the focus of theoretical studies. Moreover, a continuum two-phase model for hydrogel sheets, both for neo-Hooke and Gent-type rheology, is also new and has a multitude of applications, in particular in biology, such as concerning the morphological shapes and transitions of cells and tissue [54–56]. We thus focus a comprehensive analysis of the class of hydrogel sheets and its applications in a companion paper.

### Model parameters and forces

The geometry of the hydrogel sheet is fixed as described in Fig. 2 with *L* = 1 so that 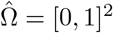. In order to measure the mechanical response of the hydrogel or hydrogel-ABM system we apply an inhomogeneous Dirichlet boundary condition (i.e. an external tensile load) for the deformation of the form

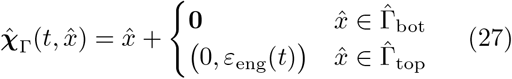

with engineering strain *ε*_eng_(*t*) = *ε*_max_ min(*t*, 1), i.e. the material is stretched for 0 ≤ *t* ≤ 1 and kept at a constant strain *ε*_eng_ = *ε*_max_ for *t >* 1. The engineering strain is related to the length of the deformed domain *L* + *δL* via *ε*_eng_ = *δL/L*. Note that depending on the material law, certain elastic materials can only sustain a certain maximal engineering strain at which the stress or the deformation become singular. Since the bulk modulus does not depend on the concentration, we can rescale the free energy density of the system such that *μ* = 1. Using typical dimensional cell traction forces of 10^−8^ *N*, typical hydrogel geometries with *HL* = 10^−8^ … 10^−6^ *m*^2^ and elastic moduli of soft tissues *μ* = 10^3^ … 10^4^ Pa, we obtain a range of nondimensional traction forces of *f*_trac_ = 10^−8^*N/*(*μHL*) = 10^−6^ … 10^−3^, e.g. cf. [5, 57]. In order to enhance nonlinear effects, we will use *f*_trac_ = 10^−3^.

The dynamic model has different time scales: The intrinsic time scale for the ABM motility and orientation dynamics is given by the time step size *τ*, as cells are updated each time step. However, the practical migration rate then also depends on the magnitude of differences in the deformation function generated by cell-cell and cell-ECM interaction. The typical diffusive and reactive relaxation time of the hydrogel is encoded in the mobilities *m* and *n* in (20). For the mobility of the hydrogel we consider two limiting cases, where either *m* = *n* = 0, so that no diffusion or solvent transport occurs and the response of the hydrogel is *purely elastic*. The other limiting cases are either of *conserved* Cahn-Hilliard type, i.e. if *m* = 10^5^, *n* = 0, or of mixed *non-conserved* Allen-Cahn type, i.e. if *m* = *n* = 10^5^. For the Cahn-Hilliard evolution this implies that after each time step with *τ* = 10^−3^ the solvent concentration is practically in equilibrium with constant chemical potential *η* and 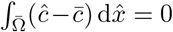 for the given applied strain. The viscoelastic relaxation time is assumed to be smaller than the time step size and thus the evolution is dominated by the ABM’s cell dynamics.

For the Allen-Cahn evolution this equilibrium assumption implies that at each time *t* the chemical potential is close to zero *η*≈ 0 and the free energy is minimal under the given applied strain, including the possibility of solvent exchange with the surrounding. For the Gent model we use a moderate limiting deformation *J*_*m*_ = 2, consistent with values for biological materials discussed in Sec. 2.1. The prefactor *γ* in the chemical energy is chosen small *γ* = 5· 10^−4^ to regularize the diffusion in regions of strong convection in regions of large stress differences, e.g. near corner singularities of the elastic sheet. We use representative parameters *α* = 4 and 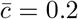 and systematically study the impact of different hydrogel energies with *k*_1_ = *k*_2_ = *k*. We mentioned before, in order to ensure that initially the hydrogel is in equilibrium, we need to set 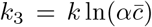, where by definition 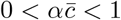.

In the following, we discuss the ingredients to compute the mechanical stress for an effective 2D hydrogel sheet for any given applied (engineering) strain *ε*_eng_. For the three-dimensional hyperelastic material with energy density *W* (***F***, ***c***, ∇***c***) as introduced in (1), the first Piola-Kirchhoff tensor defined in (8a) receives its main contribution from **P**_mech_ = *∂*_***F***_ *W*_mech_, which for a neo-Hookean material is

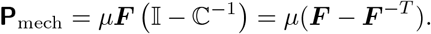

Similarly, for the Gent model (13) one has

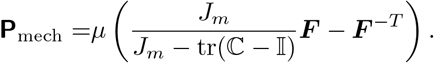

At each point these are 3 ×3 tensors (matrices). The corresponding mechanical contribution to the (symmetric) Cauchy stress is given by ***σ***_mech_ = *J* ^−1^**P**_mech_ ***F*** ^*T*^. Due to the Lagrange multiplier, we can replace *J* in *W*_chem_ by 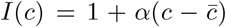, so that *W*_chem_ does not directly contribute to the mechanical stresses but only indirectly via the Lagrange multiplier that enforces the incompressibility, i.e. **P** = **P**_mech_ + *pJ* ***F*** ^−*T*^.

In the two-dimensional sheet approximation we replace the energy density by 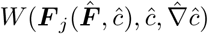, which induces some minor modifications in the effective stress tensor 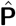. In the plane stress approximation we use the chain rule to obtain

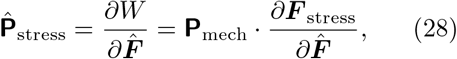

whereas in the plane strain approximation

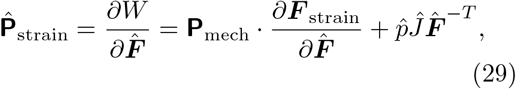

using the different expressions for 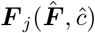 for *j* ∈ {strain, stress}, respectively. Therefore, for the thin two-dimensional hydrogel sheet, the engineering stress *σ*_eng_ is calculated in terms of the first Piola-Kirchhoff stress tensor 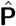 and using the normal vector 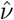 on 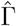 with

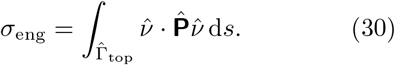

Due to the direct dependence of *W*_chem_ on *J*, due to the direct coupling with *I*(***c***) and due to the indirect coupling with the Lagrange multiplier the engineering stress of the sheet depends on all the hydrogel parameters. The same coupling will result in an inhomogeneous solvent concentration *c* when an external stress is applied.

### Purely elastic sheets

In Fig. 6, as reference curves we show the stress-strain relation for a purely elastic Gent (black lines) and neo-Hooke model (grey lines) in the plane strain (dashed lines) as well in the plane stress (solid lines) approximation. The plane stress approximation is suitable for systems of thin incompressible elastic sheets that expand and shrink in the sheet direction and change thickness without considerable bending or wrinkling. The plane strain approximation is suitable for rather thick elastic sheets, where the deformation is primarily within one plane and negligible in the orthogonal direction.

**Fig. 6.**
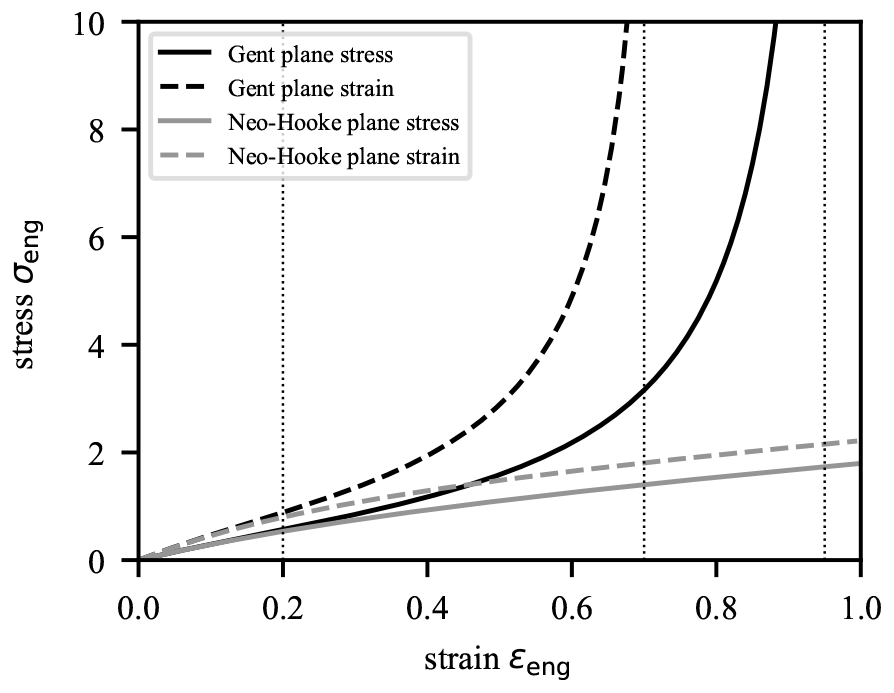
Stress-strain relation for pure elastic Gent model and pure elastic neo-Hooke model in the plane strain and plane stress approximation. The thin dotted lines show the strains for the plane stress approximation used in Fig. 7.

The strain-stiffening capability of the Gent model for both plane strain and plane stress can be directly observed in the stress-strain relation in Fig. 6, where the stress for the Gent model grows super-linearly (stiffening) and for the Neo-Hooke model it grows sub-linearly (softening). For small engineering strains 0 *< ε*_eng_ *<* 0.2, Gent and neo-Hookean materials follow the same (linear elastic) behavior with different effective elastic bulk modulus for plane strain (stiffer) and plane stress (softer), respectively. The same behavior can also be observed in Fig. 7, where we show the trace tr(ℂ− 𝕀) indicating that value of that term in the free energy density of the Gent and neo-Hookean material. For *ε*_eng_ = 0.2 both materials show qualitatively the same behavior with large deformation gradients in the corner of the elastic sheet. For larger strain *ε*_eng_ = 0.7 the strains starts to differ both in the corner and in the center of the sheet, where the neo-Hookean material has systematically larger deformations encoded in tr(ℂ − 𝕀). Near the stress singularity *ε*_eng_ = 0.95, the Gent material stiffens with tr(ℂ − 𝕀) close to *J*_*m*_, whereas the softer neo-Hookean material develops even more singular deformation gradients in the corner of the elastic sheet. In general, one may stretch the neo-Hookean sheets much further than sheets of Gent-type, since for the latter 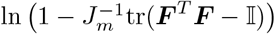 becomes singular for a certain maximal sustainable engineering strain at which the corresponding stress becomes infinite.

**Fig. 7.**
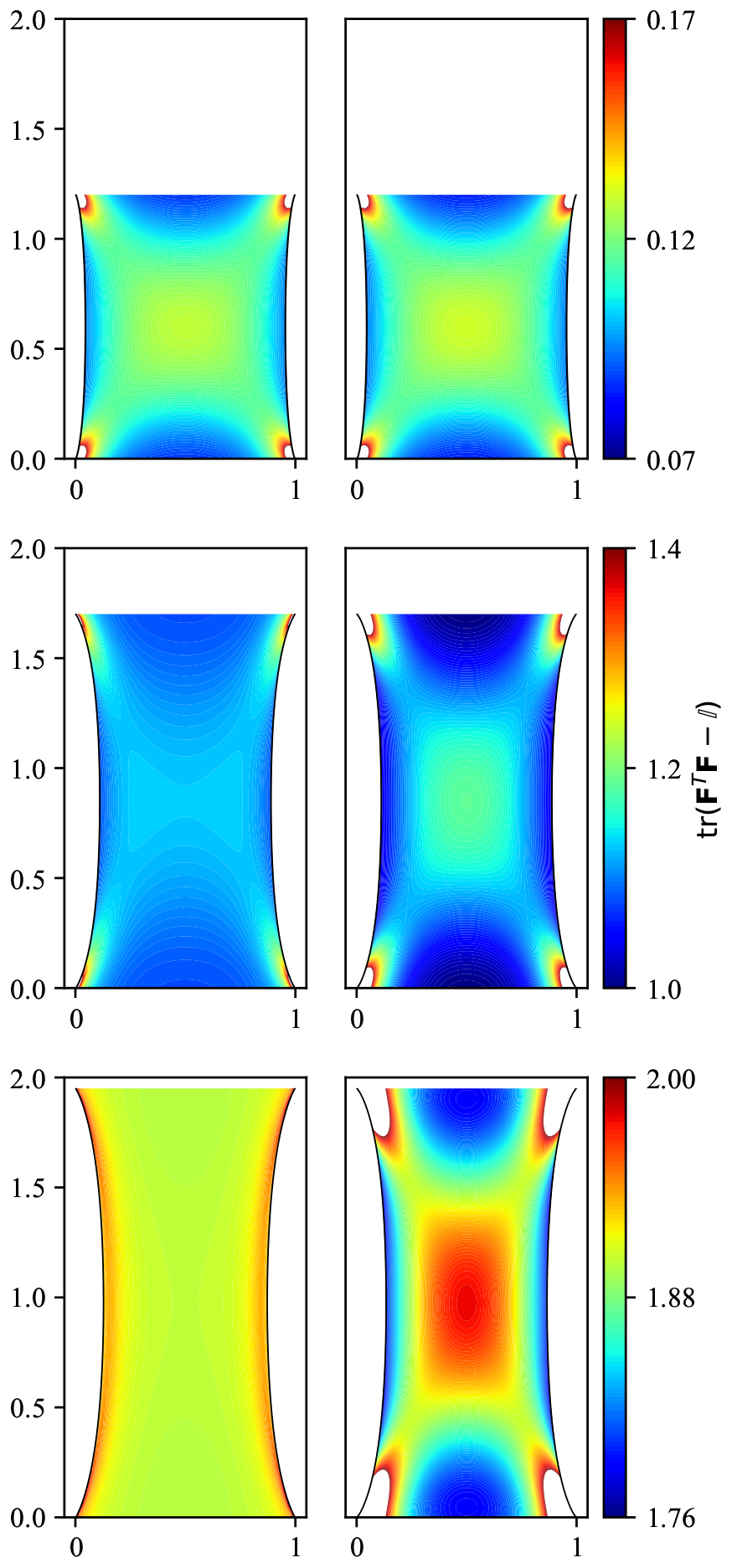
Stretching of (**left column**) pure elastic Gent material with *J*_*m*_ = 2 and of (**right column**) neo-Hookean material for increasing engineering strain (**top to bottom**) of *ε*_eng_ = 0.2, 0.7, 0.95. The colored shading in each row shows tr(***F*** ^*T*^ ***F*** − 𝕀) and for the Gent model never exceeds *J*_*m*_. In each row the (**left**) and (**right**) panel share a color bar, where white regions indicate that values exceed its range.

That maximal engineering strain depends on several parameters, e.g. on the geometry, on the elastic parameters *μ* and *J*_*m*_, but also on the hydrogel energy and the type of 2D approximation.

While strain softening also occurs in some biological tissue [8], most biological gels show strain stiffening properties. A microscopic picture is discussed in [24]. Furthermore, hydrogels also show drying induced stiffening and softening that would need to be considered additionally [45]. In this study we discuss the contribution that the hydrogel character has on stress-strain relations. While we also point out the differences between plane stress and plane strain approximation, we will focus on thin sheets best described using the plane stress.

### Hydrogel sheets

For hydrogels with conserved solvent concentration, as described by the Cahn-Hilliard equation, the stress-strain relationship of thin sheets as described by the plane stress approximation only shows small differences to that of a purely elastic material visible from the result that all full lines in Fig. 8 (top) for hydrogels for different values of *k* and the pure elastic material overlap. This behavior is shown for Gent-type materials and is similar for neo-Hookean hydrogels (not shown). However, in the plane strain approximation this changes. In this case the stress-strain curves are dependent on the values of the parameter *k* in the Flory-Huggins energy, i.e for large values of *k* the stress-strain converge to the relation of the purely elastic model whereas for lower *k* values the engineering stress reduces, see the dashed lines in Fig. 8 (top). Regarding the independence of the stress-strain relation for the plane stress case we note that the mechanical energy is expanded around the reference state with ***F*** = 𝕀 and 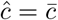 so that variations are smaller compared to a purely elastic model for a hydrogel expanded around the dry state *ĉ* = 0 as the reference. Again, this behavior is qualitatively similar for a neo-Hookean material (not shown).

**Fig. 8.**
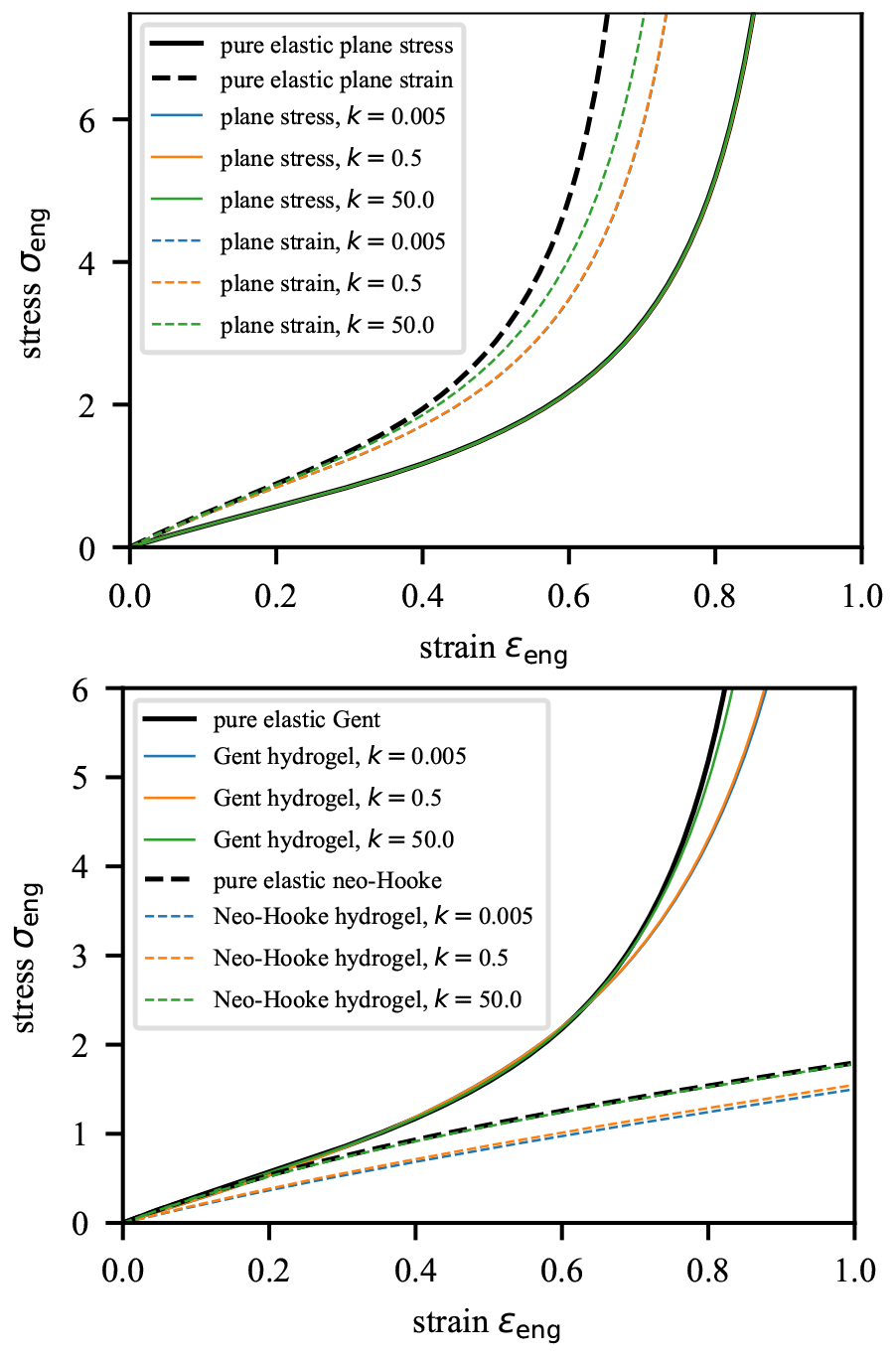
**(Top)** Stress-strain relations for Gent-type hydrogels with plane stress and plane strain for different hydrogel parameters *k* compared to pure elastic behavior (vanishing mobility *m* = *n* = 0). Note that *all* full lines overlap. **(Bottom)** Stress-strain curves for thin sheets (plane stress): purely elastic Gent model vs. Gent hydrogel model with the Allen-Cahn equation. Hydrogels are considered for different parameters *k*_1_ = *k*_2_ = *k* compared to pure elastic models.

However, if we allow for a solvent flux into or out of the hydrogel to describe exchange with a surrounding solvent bath, then also in the plane stress limit the stress-strain relation and the materials stiffness differs visibly from that of a pure elastic material, cf. Fig. 8 (bottom). For large constants *k*, the material behavior is closer to that of the pure elastic material and for smaller *k* the material gets softer. This effect should not be confused with drying induced softening or stiffening but is purely related to the change in hydrogel volume that allows a further reduction of the total free energy of the system as opposed to the incompressible (conserved) elastic system, where such a change of volume is impossible. These observations demonstrate the importance of distinguishing between stiffening and softening effects in hydrogels that are caused by strain, solvent drying or even just solvent redistribution by diffusion. The main difference between the stress-strain relation of the neo-Hookean hydrogel and the Gent hydrogel visible in Fig. 8 (bottom) is that while for neo-Hookean hydrogel the deviation from pure elastic behavior is evident for all strains, for Gent-type hydrogels it is only clearly visible for large strains.

In Fig. 9 first row, tr(***F*** ^*T*^ ***F*** − 𝕀) at an engineering strain of *ε*_eng_ = 0.94 for both models, the neo-Hooke and Gent hydrogel with with conserved solvent concentration and non-conserved solvent concentration for different *k* is presented. Furthermore, the corresponding solvent concentrations are highlighted int Fig. 9 second row.

**Fig. 9.**
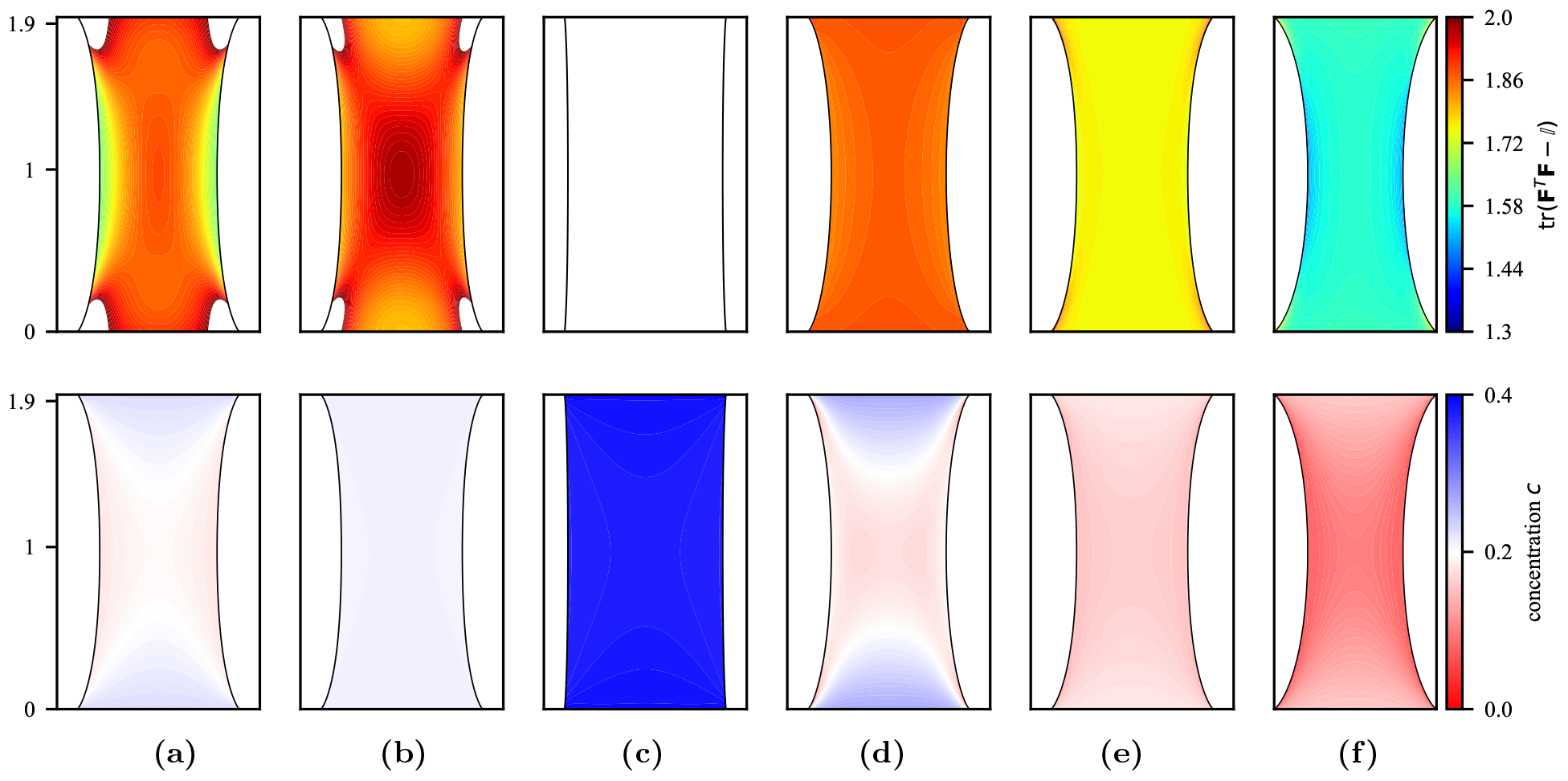
Comparison of stretched hydrogels in the plane stress approximation at an engineering strain *ε*_eng_ = 0.94. In the first row tr(***F*** ^*T*^ ***F*** − 𝕀) is illustrated, while in the second row the solvent concentration *c*. From left to right different hydrogels and configurations are presented: **(a)** neo-Hooke hydrogel with Cahn-Hilliard evolution and *k* = 0.5, neo-Hooke hydrogel with Allen-Cahn evolution and **(b)** *k* = 50 or **(c)** *k* = 0.5, **(d)** Gent hydrogel with Cahn-Hilliard evolution and *k* = 0.5, Gent hydrogel with Allen-Cahn evolution and **(e)** *k* = 50 or **(f)** *k* = 0.5.

Interestingly, Fig. 9 reveals significant qualitative differences between neo-Hookean hydrogels and Gent-type hydrogels upon stretching. Firstly, while for large *k* both hydrogels behave like their elastic counterparts, for sufficiently small *k* the neo-Hookean hydrogel absorbs solvent upon stretching and therefore, increases its volume, the Gent-type hydrogel releases solvent upon stretching and therefore, reduces its volume. This effect can be most prominently seen in the lower row of Fig. 9 in c) showing absorption and in f) showing release of solvent. In particular for the neo-Hookean material this results in the strikingly different cross-section shape, where the deformed side boundaries are almost straight due to solvent absorption. With conserved solvent concentration, the lower panels a) and d) show that solvent diffusion due to stretching leads to lower solvent concentration in the middle area of the sheet and higher solvent concentration near the top and bottom boundaries. Additionally, in the upper panel d) we observe that strain-stiffening effect already observed in Fig. 7 for the purely elastic material. In particular in the upper panels e) and f) one can see that with lower *k* the hydrogel mechanical energy is reduced and therefore, the material becomes softer.

The consistent coupling of such effects in biological hydrogels is a necessity for a fundamental understanding of such a complex system in order to make meaningful predictions that are able to distinguish from more complex biological processes, where cell movement depends on biochemical signaling processes in addition to mechanical stimuli, which can be modeled with such a system.

## 4 Collective migration of cells on hydrogel sheets

The main focus of this section is the discussion of interactions between cells and hydrogels, i.e. with Gent-type or neo-Hookean elastic models under the plane strain or plane stress condition for conserved (Cahn-Hilliard) or non-conserved (Allen-Cahn) solvent concentration. We concentrate on choices that have a clear impact on the resulting cell migration and pattern formation. In addition, we will investigate the interaction of cells with purely elastic materials under the plane strain and plane stress condition to complete our study. Since for the unstretched situation, Gent type and neo-Hooke materials give comparable results, since we are essentially in linear regime, we will discuss our results only for the purely elastic neo-Hooke model.

Notice that we will use for all simulations the same (random) initial distribution of 500 cells for better comparison of the pattern formation. Also, note that due to the different boundary conditions on top/bottom and the side boundaries, cells can have a preferred orientation even without any stretching of the sheet. This orientation may change once we apply a (engineering) strain *ε*_eng_ by stretching the sheet. We will consider this in Sec. 4.1. We then investigate the effects on cell migration and pattern formation for vanishing strain *ε*_eng_ = 0 in Sec. 4.2. In Sec. 4.3 we provide a qualitative analysis of the phenomena observed by detailed local case studies of the deformation function used by the ABM.

The evolution of cell patterns and the cell’s orientations is accompanied by the corresponding evolution of the number of horizontal, vertical and diagonal cells, where we define the effective dipole count via

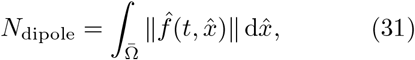

which reduces if the dipole force field of adjacent cells overlap fully (same orientation) or partially (different orientation) and therefore serves as an effective count of cell chains.

In the upper left and right panel of Fig. 10 we observe that for early times 0 *< t <* 0.1 the neo-Hookean and Gent-type hydrogel evolve the same, since they have the same linear material behavior. Remarkably, between *t* = 0.05 and *t* = 0.1 we can observe the change from mainly vertical to mainly horizontal orientation of cells. Usually, the response of cell orientation due to a change of local elastic behavior is relatively fast compared to migration and chain formation of cells. Diagonal orientation of cells is negligible for both neo-Hookean and Gent-type materials for most of the time. This effect is caused by the non-isotropic nature of dipole positions that for the moment need to be aligned with the computational mesh and should be overcome by future ABM that allow generic cell orientations. Currently, only strongly favorable diagonal strains will lead to diagonal cell orientations. In the lower panels of Fig. 10 the neo-Hookean horizontal orientation for 1 *< t <* 3 stays basically constant, while for the Gent-type hydrogel the number of horizontal cells reduces at the expense of vertical and diagonal cells that accumulate at the top and bottom boundary. Considering the effective dipole number, cell chains continue to grow in length for the neo-Hookean material, while for the Gent-type material, the rapid change in cell configuration during the interval of 0.5 *< t <* 1 leads to a reduction in chain length.

**Fig. 10.**
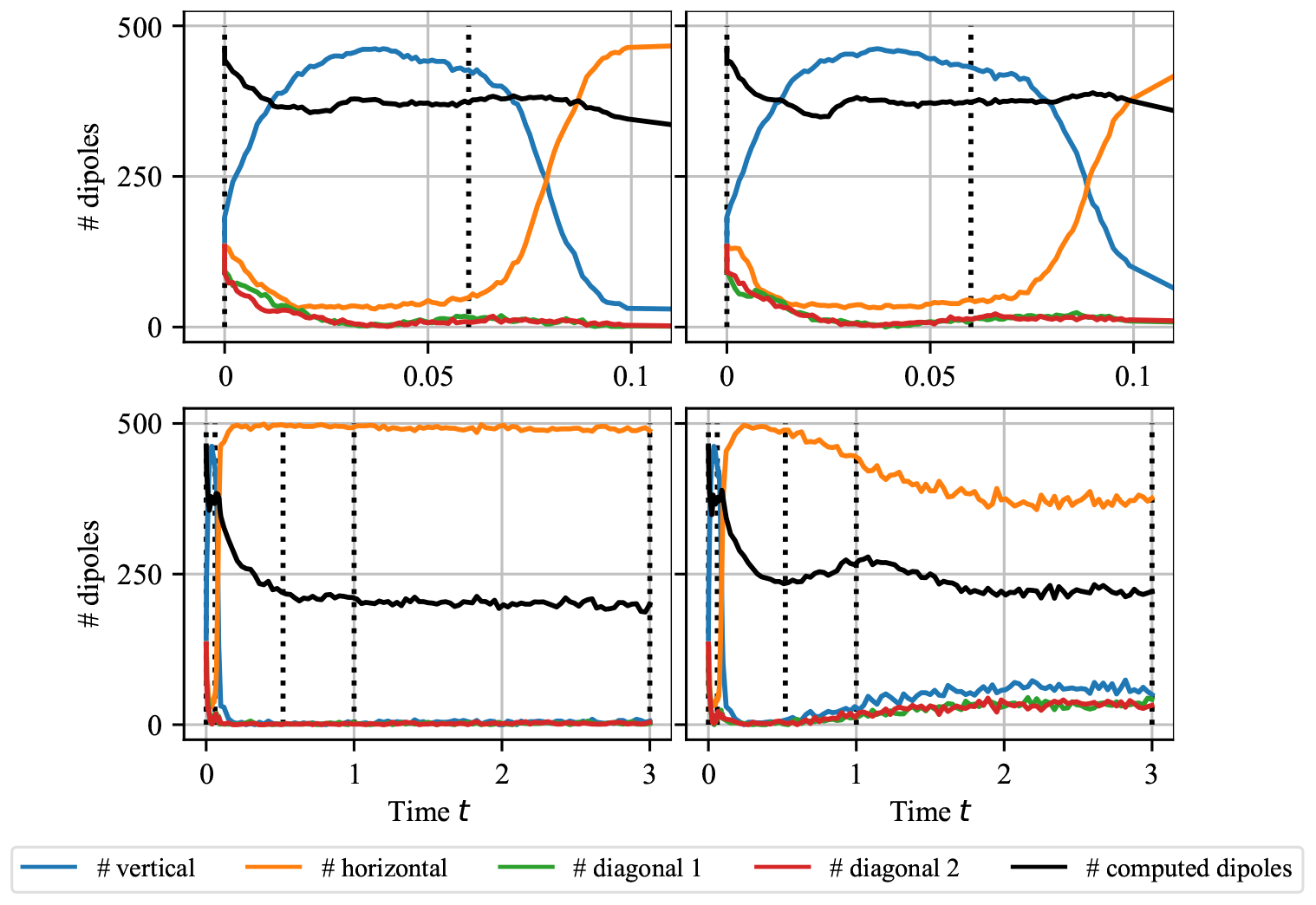
Number of cells with horizontal, vertical or diagonal orientation and effective number of computed dipoles in a stretched hydrogel with *ε*_eng_ = *ε*_max_ min(*t*, 1) with *ε*_max_ = 0.9 computed **(left)** for the neo-Hookean hydrogel and **(right)** for the Gent-type hydrogel. In the (**lower panel**) the history 0 ≤ *t* ≤ 3 and the the (**upper panel**) the initial stage for 0 ≤ *t* ≤ 0.1 is shown. The dotted lines indicate the time of the snapshots shown in Fig. 11 and Fig. 12, i.e. *ε*_eng_ = *{*0, 0.06, 0.51, 0.9, 0.9*}* from left to right.

### 4.1 Cells on stretched sheets

In accordance with Sec. 3, we begin the analysis of the interaction of cells and hydrogels by analyzing the patterning, orientation and migration of cells for stretched hydrogels. For this, we first consider the evolution of hydrogels with non-conserved solvent concentration in the plane-stress limit for neo-Hooke and Gent-type materials. We consistently use *n*_cells_ = 500 cells with a traction force *f*_trac_ = 10^−3^ for a single cell, which is a typical value given in the literature, e.g. in [5]. For cell traction forces on this scale, effects of nonlinear elasticity will be negligible. Here, we investigate the cells’ response by applying a finite strain via stretching the hydrogel sheets in our numerical stretching experiments.

Fig. 11 shows the typical evolution of strain and solvent concentration and the corresponding cell patterns during the initial stretching 0≤ *t* ≤1 of a neo-Hookean hydrogel **(a-d)** and for later times **(e)** *t* = 3. It can be seen that the initially random cell configuration from panel **(a)** at *t* = 0 leads to a vertical orientation of the cells after a short time, as can be seen in **(b)**. Beyond a strain of *ε*_eng_ = 0.1, the cells orient themselves horizontally **(c)** and after some time form longer chains **(d)**. As the chain formation progresses further, on a longer time scale cells accumulate at the lower and upper boundary of the thin hydrogel sheet, **(e)** at *t* = 3. However, since the neo-Hookean hydrogel does not contract as much due to solvent absorption as previously observed in Fig. 9**(c)**, a significant proportion of the cells remain in the inner area of the sheet.

**Fig. 11.**
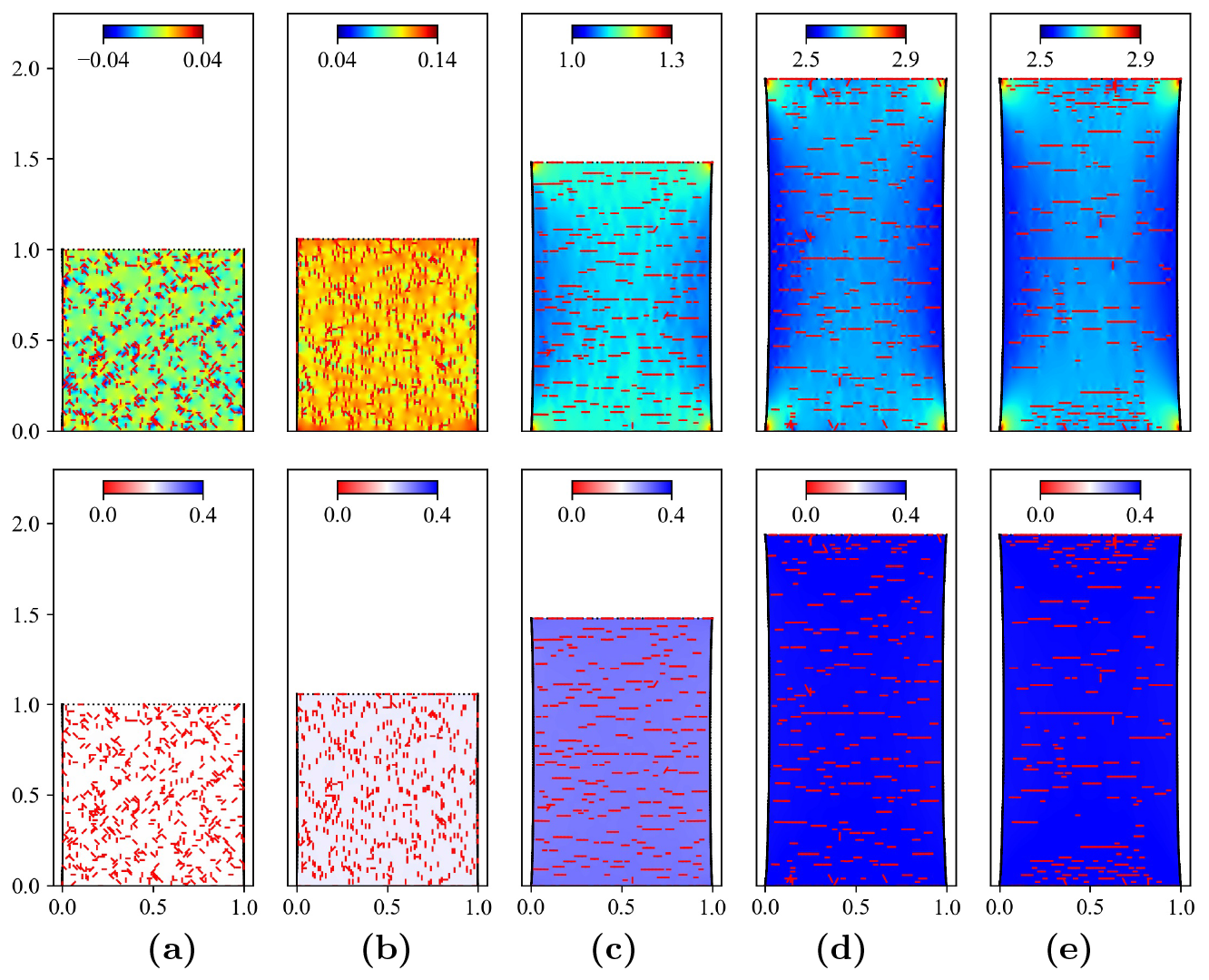
Neo-Hookean hydrogel with *k* = 0.5 and non-conserved solvent concentration for increasing strain 0 ≤ *ε*_eng_ ≤ 0.95 from left to right showing **(a)** initial configuration, **(b)** *ε*_eng_ = 0.06, **(c)** *ε*_eng_ = 0.51 and **(d**,**e)** *ε*_eng_ = 0.9. The (**upper panel**) shows strains tr(***F*** ^*T*^ ***F*** − 𝕀) (Note: color bar changes) and the (**lower panel**) shows concentrations *ĉ*.

From the point of view of the loading and hydrogel parameters, the stretching experiment with the Gent-type model in Fig. 12 **(a-c)** is performed analogously. Only, as already shown in Fig. 9, the solvent leaves the Gent-type hydrogel and thus reduces the total volume. While the same flipping from vertical to horizontal cell orientation can be observed as for the neo-Hookean material, especially at the beginning of the tensile experiment, the much stronger constriction of the hydrogel leads to a stronger migration of the cells and thus to a rapid accumulation at the lower and upper edge of the thin layer visible in **(d)** and **(e)**. The horizontal orientation of cells for the highly stretched Gent-type material is experimentally unexpected as for the stiffening material alignment with the direction of the stretch would be expected, cf. [13]. Our model for the hydrogel sheet predicts horizontal alignment, which is caused by the definition of the deformation function preferring a horizontal alignment since for sufficiently large external stretch |*F*_*xx*_−1| is smaller than |*F*_*yy*_− 1| for both neo-Hookean and Gent type materials. However, biological cells constantly reattach to the ECM and therefore only measure changes in the deformation relative to an external field, i.e. the stiffness being the second derivative of the energy density. This property of the cell-ECM interaction might suggest an extension of the ABM for external strains.

**Fig. 12.**
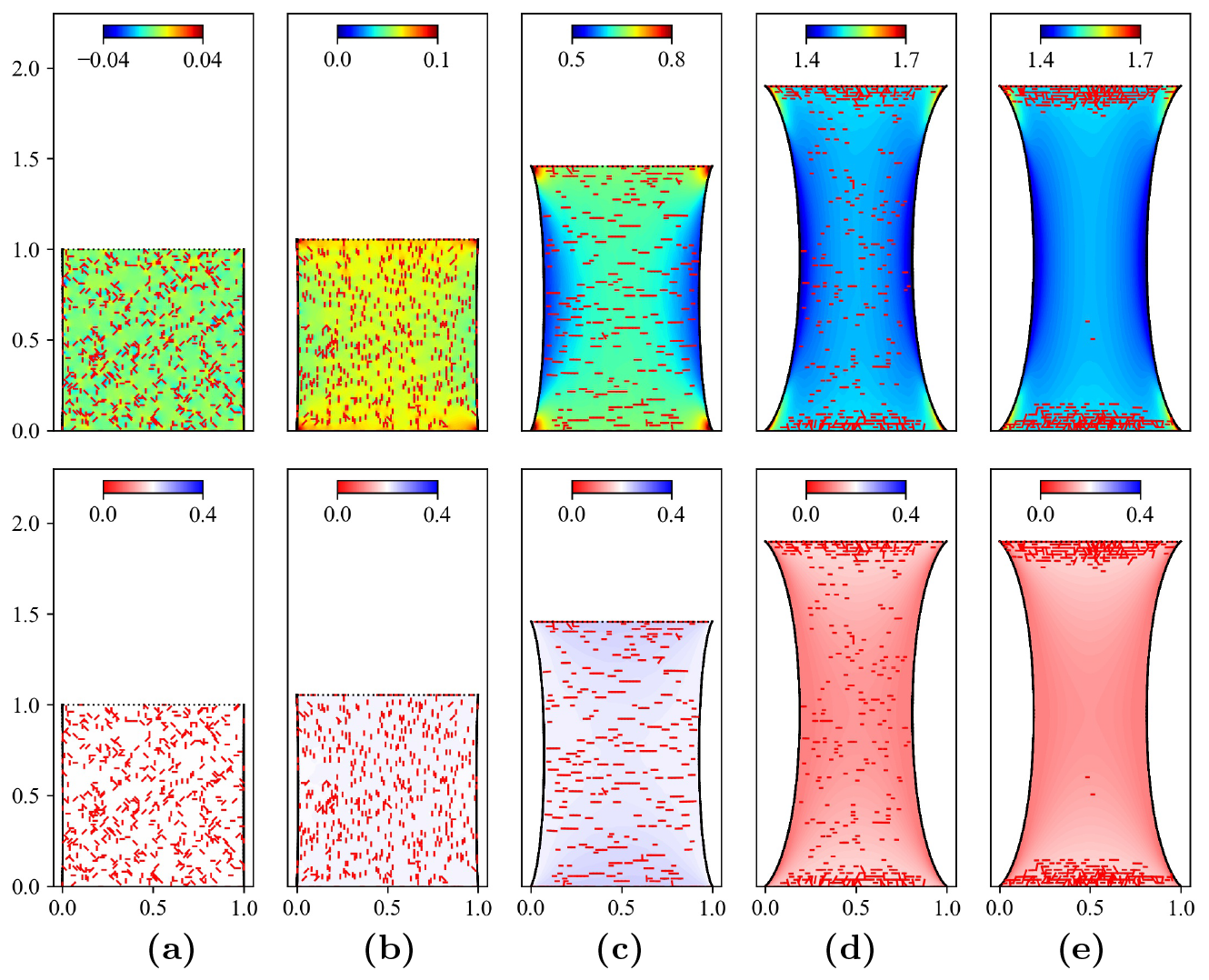
Gent-type hydrogel with *k* = 0.5 with non-conserved solvent concentration and increasing strain 0 ≤ *ε*_eng_ ≤ 0.95 from left to right for **(a)** initial configuration, **(b)** *ε*_eng_ = 0.06, **(c)** *ε*_eng_ = 0.51 and **(d**,**e)** *ε*_eng_ = 0.9. The (**upper panel**) shows strains tr(***F*** ^*T*^ ***F*** − 𝕀) and the (**lower panel**) shows concentrations *ĉ*.

In the later Sec. 4.3 we provide a qualitative explanation for the different orientation and the cell migration due to the cell-hydrogel interaction and for the chain formation due to the cell-cell interaction.

### 4.2 Cells on relaxed sheets

Our next goal is to investigate the cell interaction with the unstretched materials, starting with purely elastic models. As before, we always start with the same initial cell configuration to allow for a better comparison of the cell dynamics. Furthermore, for unstretched elastic sheets and small traction force by the cells, Gent and neo-Hookean materials behave quite similar, as they are still in their linearly elastic regime. We thus focus our analysis here only on the neo-Hookean model.

We begin with a comparison of the number of vertical, horizontal and diagonal dipoles. Fig. 13 shows that the number of diagonal dipoles decreases rapidly and the cells are mainly horizontally and vertically orientated.

**Fig. 13.**
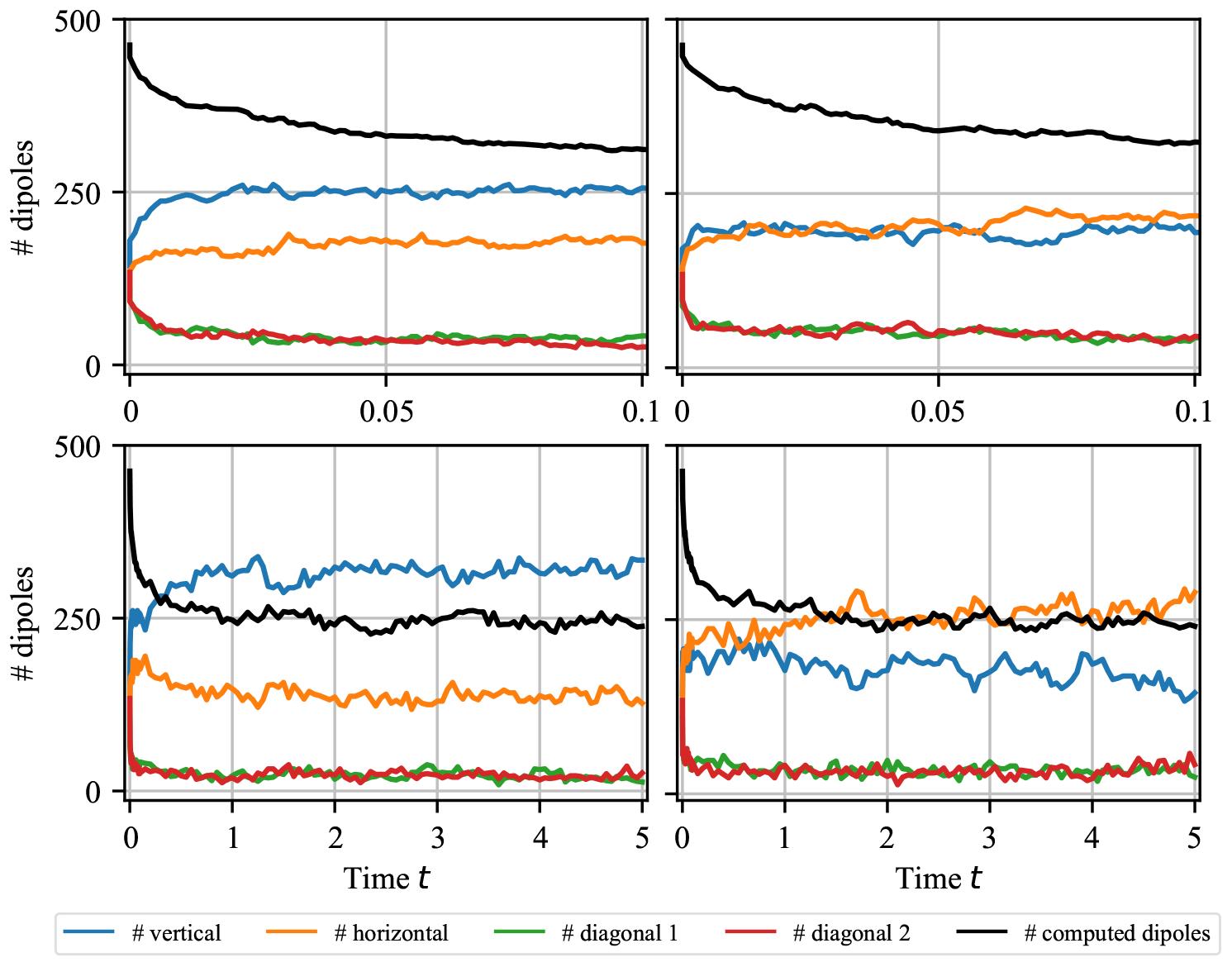
Number of cells with different orientation and effective dipole count for the unstretched pure elastic neo-Hookean material **(left)** under plane stress condition and **(right)** under plane strain condition.

In the plane strain case shown in Fig. 13 **(right)** there is no clear preference for one orientation compared to the plane stress case shown in Fig. 13 **(left)**, where the cells favor a vertical orientation. The effect in the plane stress situation has been observed before in [5]. Interestingly, in both situations the number of computed dipoles is comparable. This indicates that the cells form long-range alignments in both cases.

Considering the long-time cell patterns Fig. 14 illustrates that under the plane strain condition cells tend to aquire a more horizontal orientation. In the plane strain situation almost the complete fixed boundaries are covered by cells. Although cells have a clear preference to stay at positions with small deformation in the plane stress case, fewer number of cells move to the clamped boundaries. Furthermore, in the plane stress case the cells form long-range alignments in the vertical direction, while in the plane strain case the alignments are shorter and a clear vertical or horizontal orientation is not present. If one compares the temporal development of the dipole orientations in Fig. 13 in combination with the cell distribution of the clamped boundary, it can be seen that the cells away from the clamped boundary are almost uniformly oriented vertically and horizontally under the plane strain condition, while the cells are quickly oriented vertically under the plane stress condition. This means that in the plane stress case one has less ‘active dipoles’ in the sense that aligned cells applying traction forces to the same node cancel their forces out and one has a net force of zero at this node. In addition, in areas where one has a high concentration of cells forming long-range alignments with the same orientation the elastic material experience almost no deformation. If the cells on the clamped boundary are ignored, the difference between vertical and horizontal dipoles would increase in the case of plane stress, while in the case of plane strain, the number of vertical and horizontal dipoles would become much closer.

**Fig. 14.**
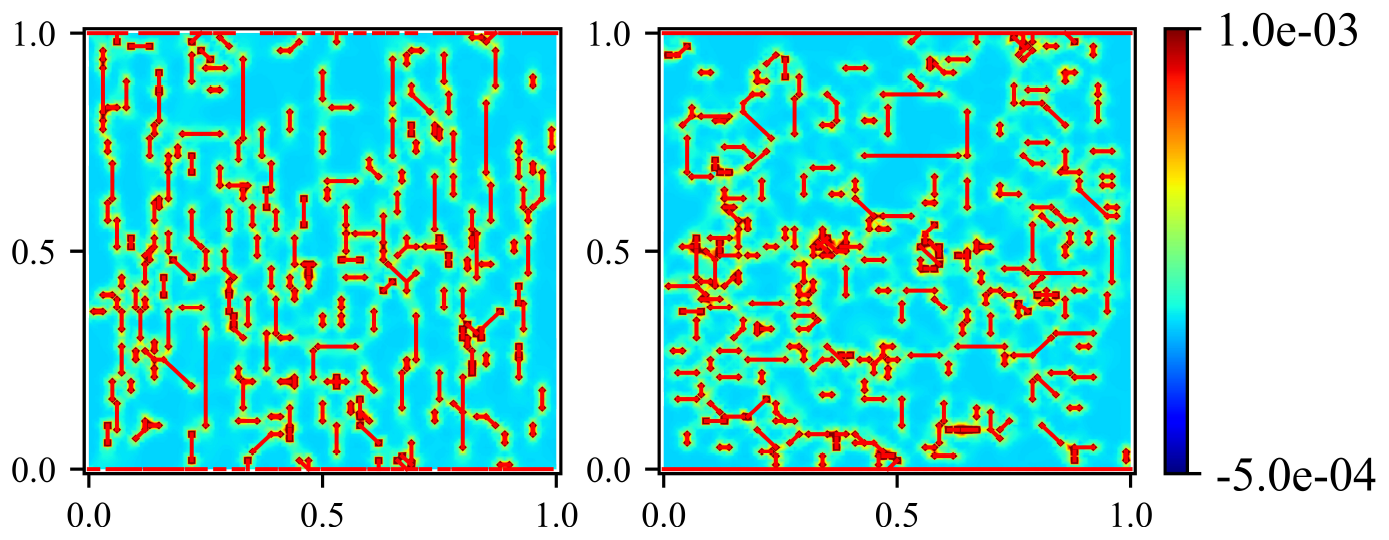
Final cell configuration without stretching after *T* = 5, i.e. 5000 iterations. Purely elastic neo-Hookean material **(left)** under plane stress condition and **(right)** under plane strain condition. The colored surface represents strain tr(***F*** ^*T*^ ***F***− 𝕀) and the red lines the cells.

A comparison of the corresponding hydrogels under plane strain and plane stress condition shows similar results and we focus here on hydrogels under plane stress. In addition we briefly compare the neo-Hooke hydrogel for the Allen-Cahn (non-conserved) case, the Gent-type hydrogel for Allen-Cahn case and the Gent-type hydrogel for the Cahn-Hilliard (conserved) case, for *k* = 0.5. We obtain comparable behavior for unstretched and stretched hydrogels for the Cahn-Hilliard case and the corresponding purely elastic materials based on our previous findings. This is also reflected in Fig. 15 **(right)** and Fig. 16 **(right)**. Furthermore, Fig. 15 and Fig. 16 show that under plane stress the hydrogels behave quite similar due to the relative small deformation, which we already highlighted in Sec. 4.1.

**Fig. 15.**
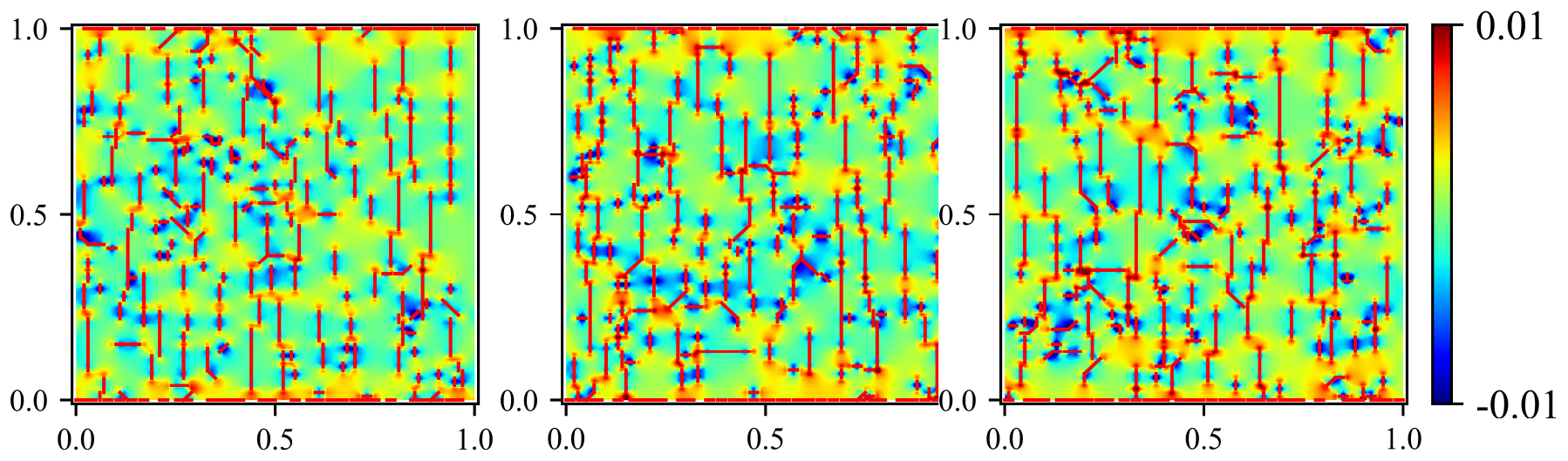
Cell configuration without stretching at *T* = 5, i.e. after 5000 iterations. Neo-Hooke hydrogel with Allen-Cahn evolution and *k* = 0.5 **(left)**, Gent hydrogel with Allen-Cahn evolution and *k* = 0.5 **(middle)**, and Gent with Cahn-Hilliard evolution and *k* = 0.5 **(right)** under plane stress condition. The colored surface represents strain tr(***F*** ^*T*^ ***F*** − 𝕀) and the red lines the cells.

**Fig. 16.**
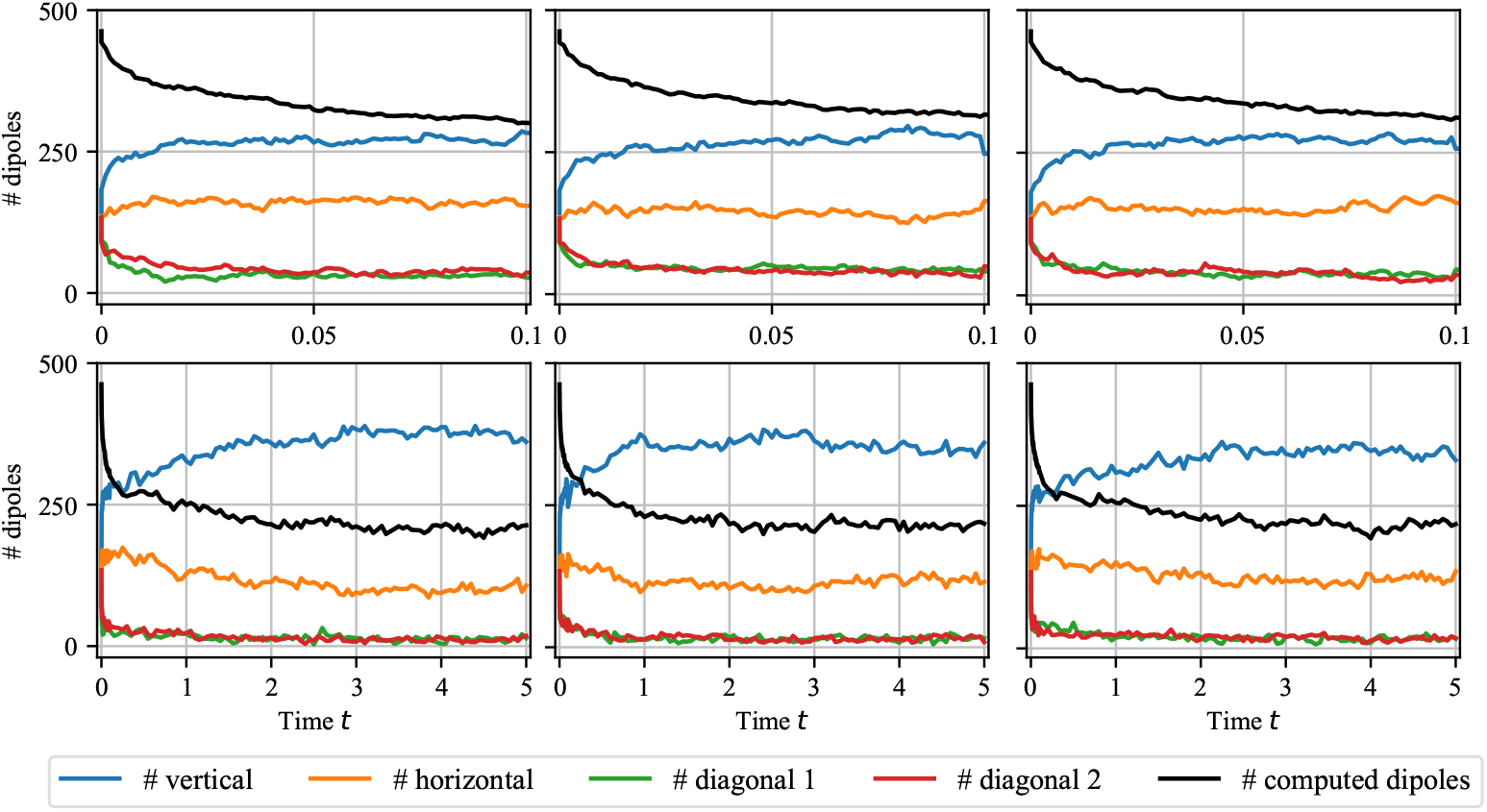
Number of cells with horizontal, vertical or diagonal orientation and effective number of computed dipoles for the neo-Hooke hydrogel with Allen-Cahn evolution and *k* = 0.5 **(left)**, for the Gent hydrogel with Allen-Cahn evolution and *k* = 0.5 **(middle)**, and for the Gent with Cahn-Hilliard evolution and *k* = 0.5 **(right)** under plane stress condition.

### 4.3 ABM for cell-cell and cell-hydrogel interactions

The simulations of the ABMs in Sec. 4.1 and Sec. 4.2 show that cells migrate and orient on both relaxed and on stretched hydrogels. Both migration and orientation are affected by the type of hydrogel model, the cell-cell and cell-hydrogel interactions, governed by the ABM via the deformation function 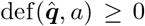 from (26). Due to the update proposed in the ABM Table 1, cells 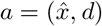favor positions 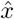 and orientations *d* for which the deformation function is minimal.

We now show that the main features of migrating cell populations on hydrogels as observed in Sec. 4.1 and Sec. 4.2, can be understood by an analysis of the hydrogel-cell system for cell configurations ***a*** ∈ ***A*** with as few as *n*_cells_ = 1 (cell-hydrogel) and *n*_cells_ = 2 (cell-cell) cells.

#### Cell-hydrogel interaction

For a given time-independent single cell state 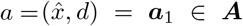 with *n*_cells_ = 1 we compute the traction force 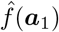 and solve the corresponding (stationary) hydrogel problem for 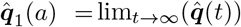. With this solution we evaluate the 1-cell deformation

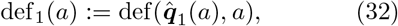

in order to evaluate, for example, the most likely orientation. Therefore, for fixed cell position 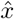 one needs to find the orientation *d* ∈ ∆ for which 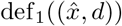 is smallest. Similarly, by plotting the function 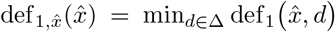 one finds the places where the deformation function has a local or global minimum and therefore are places that cells would migrate to by cell-hydrogel interaction. In Fig. 17 the corresponding comparison of the neo-Hookean elastic material in the plane strain and in the plane stress approximation is shown without additional strain, i.e. *ε*_eng_ = 0. While in the plane stress approximation vertical orientation is favored at all points, in the plane strain approximation the central region also supports horizontally oriented cells. Note that ultimately, due to the boundary condition cells would also favor locations close to the top and bottom boundary due to the smaller deformation, where the cell states are constrained to horizontal orientation. These observations also support our findings in Fig. 13, where the plane strain configuration has a higher count of horizontal cells due to cells at the center and cells at the lower and upper boundary. The trend of cells migrating to the top and boundaries was observed in Fig. 14.

**Fig. 17.**
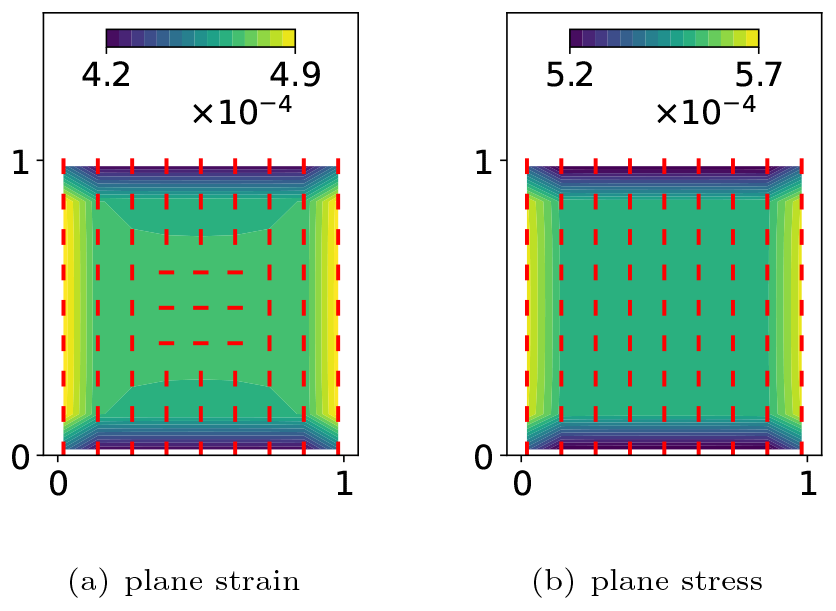
Cell-hydrogel interaction from single cell computation showing def_1_(*a*) for different neo-Hookean elastic materials. The red line indicates the optimal orientation and the colors shows the optimal cell location based on the smallest value of the deformation function.

Note that we can also quantify the transition of orientation and migration due to stretching by computing the single-cell deformation function on slightly stretched neo-Hookean sheets in Fig. 18. Here one can clearly observe that for small strains *ε*_eng_ *<* 0.1 (plane strain) and *ε <* 0.05 (plane stress) the vertical orientation is even enhanced but breaks down for large strains. In particular close to the transition, the local deformation function looks quote different in the the plane stress (*H* ≪*L*) and plane strain case (*H* ≫*L*). Observing such a strain-induced change of orientation would of course strongly support a certain strategy for cell motion and orientation encoded in an ABM. In Fig. 19 we extend the single-cell analysis to strongly stretched neo-Hookean hydrogels (right) compared to pure elastic materials (left). While both show a clear preference for a horizontal cell orientation, the variations in the single-cell deformation function are much smaller leading to a somewhat smaller migration rate.

**Fig. 18.**
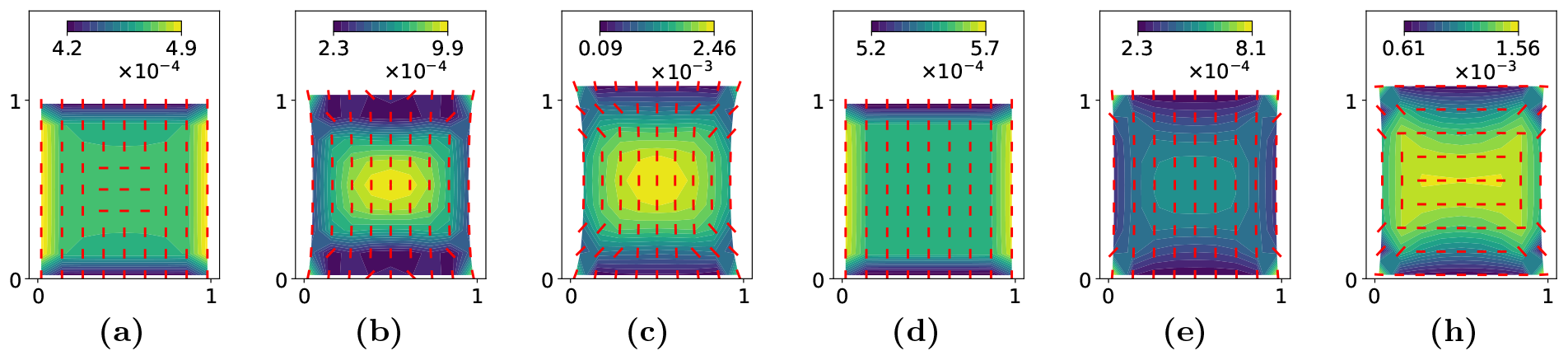
Single cell deformation def_1_(*a*) on unstretched or slightly stretched Neo-Hookean elastic sheets in **(a**,**b**,**c)** plane strain or **(d**,**e**,**f)** plane stress approximation, where **(a**,**d)** are unstretched *ε*_eng_ = 0 and **(b**,**e)** are slightly stretched *ε*_eng_ = and **(c**,**f)** are slightly more stretched *ε*_eng_ = 0.1.

**Fig. 19.**
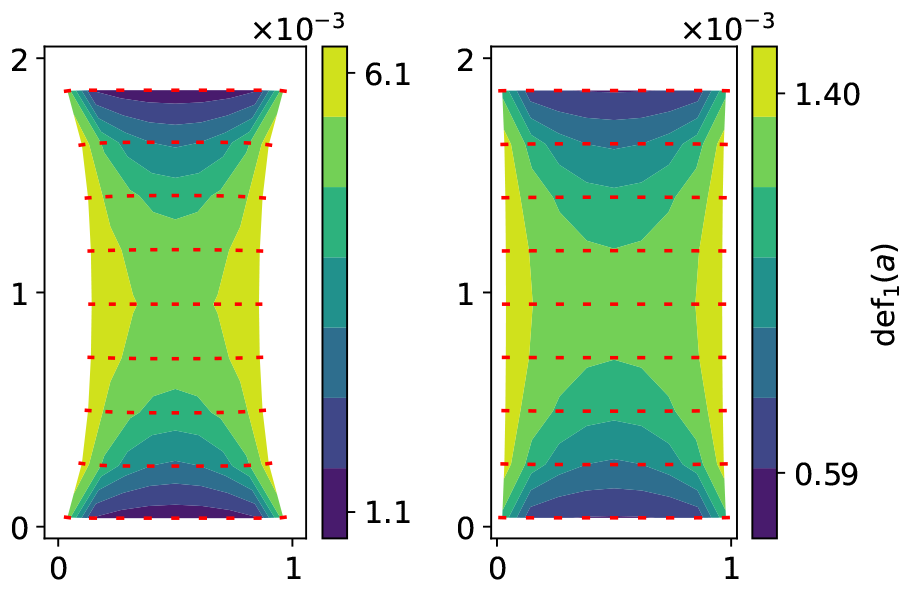
Cell-hydrogel interaction for strongly stretched elastic sheet from single cell computation showing def_1_(*a*) for neo-Hookean material in the plane stress approximation. We show **(left)** the pure elastic material and **(right)** the non-conserved solvent with *k* = 0.5

#### Cell-cell interaction

Similarly, the cell-cell interaction can be studied using two time-independent cells ***a***_2_ = (*a, a*^∗^) ∈ ***A*** with *n*_cells_ = 2. Here, we fix one cell at the center of the domain with a fixed orientation, i.e. 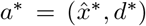 with 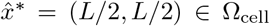 and *d*^∗^ = *d*_↕_. Then we set 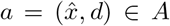 arbitrary, compute the traction force 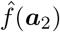 and solve the corresponding (stationary) hydrogel problem for 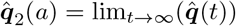. Based on this solution, we define the two 2-cell deformations

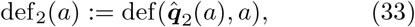

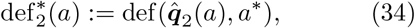

relative to the given cell *a*^∗^, where up-to discretization artifacts due to the singular nature of the traction force 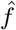, both definitions should give rise to the same interpretation of cell-cell interactions. As before, from *a* we can deduce most likely orientations and cell positions.

In Fig. 20 we show the resulting two-cell deformation function based on 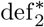 and observe that cell-cells interactions favor formation of chains with cells *a, a*^∗^ of same orientations being at positions 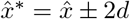. However, note that the magnitude of cell-cell interaction is ∼ 10^−4^, whereas the cell-hydrogel interaction is usually stronger with ∼ 10^−3^ and therefore dominates the cell dynamics. On the other hand, cell-hydrogel interaction is usually a long-range effect whereas cell-cell interaction is extremely short-ranged and limited to a few diameters of cell-dipole distances. The cell-cell interaction map shown in Fig. 20 confirms the previously made observation of cells forming long chains, an effect that in a transient regime might be disrupted by external strain fields as we observed in the right panel of Fig. 10. This investigation of the deformation function shows that single and double-cell investigations provide a basic understanding of the collective pattern formation and the migration and (re)orientation of cells in external strain fields.

**Fig. 20.**
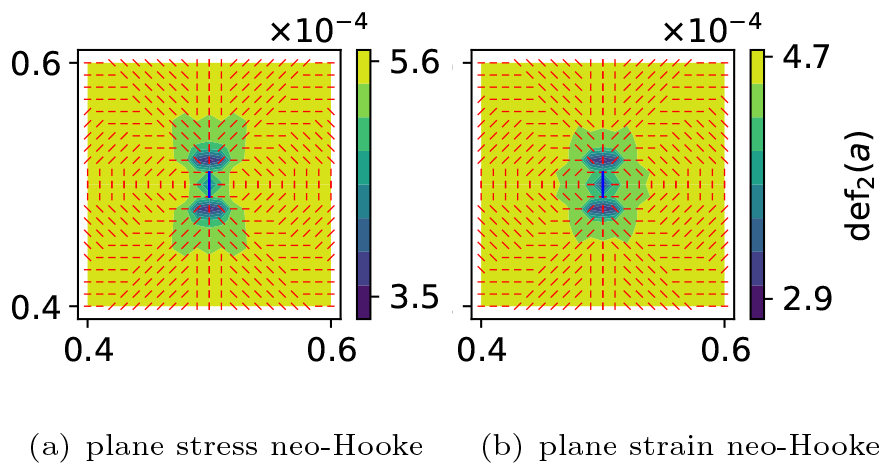
Cell-cell interaction from double cell computation showing def_2_(*a*). The red line indicates the optimal cell orientation of a second cell and the colors shows the optimal cell location based on the smallest value of the deformation function. The first cell is fixed at 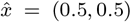 with a vertical orientation (blue line).

#### Local cell-hydrogel response

Effects of single cells on strain-stiffening hydrogels, have been observed for example in [58] or in [59, 60]. Within our current hydrogel-ABM model framework, where we use *N* = 100 for (coarse) tensorial computational meshes, such that the vertices of the triangle mesh for the finite element approximation 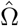 coincide with the cell positions Ω_cell_. However, since a cell at 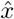 exerts a dipole-like force a neighboring mesh vertices 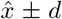, in particular the near-field of the hydrogel deformation and concentration might be massively underresolved. To see the impact, we compare in Fig. 21 deformation and concentration profile near a single cell at 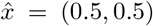 for underresolved (left panels) and highly resolved (right panels) hydrogels for neo-Hookean (upper panels) and Gent (lower panels) hydrogels. Here, instead a dipole generated by Dirac measures we approximate 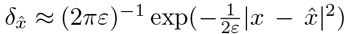 with *ε* = 10^−6^ and successively increase the traction force to its maximal value^1^ to find a stationary solution of the nonlinear hydrogel problem. This shows a good approximation of the elastic problem on a coarse grid, since for the ABM in its current form, only the deformation in the vicinity of the cell is of importance. However, in particular the effect of solvent displacement due to cell traction on the coarse grid is greatly underestimated and could have an impact on the predictive power of the fully coupled model.

**Fig. 21.**
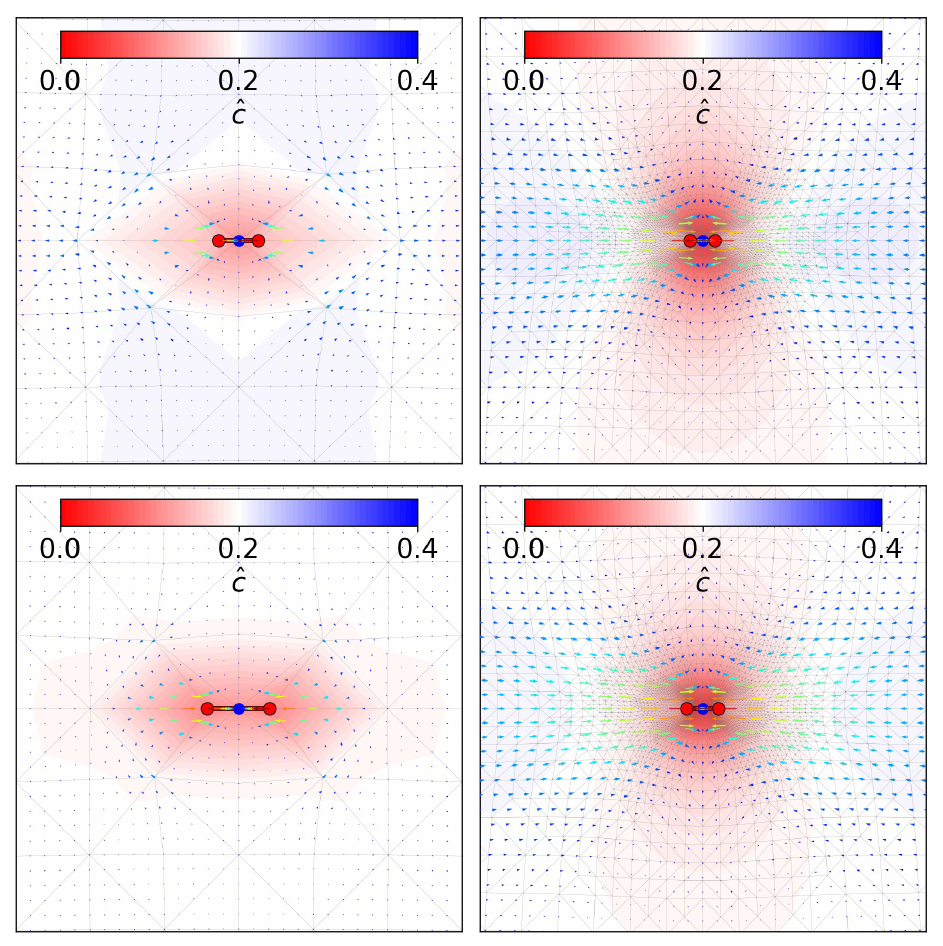
Concentration and deformation in the region 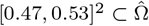 of a single horizontal cell at 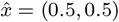 on spatially uniformly and (locally) highly resolved computational meshes. **(Upper left)** neo-Hookean uniform vs **(upper right)** neo-Hookean highly resolved and **(lower left)** Gent uniform vs **(lower right)** Gent highly resolved with large traction force *f*_trac_ = 4 10^−2^. The shading shows the concentration with red, white and blue corresponding to 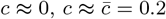 and *c*≈ 0.4. The colored vector field shows the displacement field and the thin gray mesh displays the deformed computational mesh.

In order to capture the cell’s active strain-stiffening impact on the hydrogel network higher resolutions of the numerical grid will be necessary.

## 5 Summary and outlook

We used hydrogels as a bottom-up approach for designing three-dimensional models that mimic certain properties of a native ECM environment, e.g. swelling and solvent diffusion. For this we have developed a model framework that couples a continuum two-phase model for a viscoelastic hydrogel with an agent-based model that governs the migration decisions of each cell as it interacts with the hydrogel and other cells of the population. While the coupled cell-hydrogel model framework developed in this study can in principle be used for modeling a wide range of applications in the context of cell invasion into three-dimensional hydrogel scaffolds, we focused on cell migration on scaffolds of a thin hydrogel sheet. We analyzed the geometrical set-up of a fixed hydrogel sheet for which we investigated the stress-strain state by applying specific loads at the fixed boundaries of a rectangular sheet, leaving the other two boundaries free. We compared scenarios that also allow for possible exchange of the solvent phase through the free boundaries. In addition we compared our results with those using a neo-Hooke model for the nonlinear elastic network to contrast the effects of strain-stiffening on pattern formation of the cell population. The numerical approach is based on a thermodynamic consistent mechanical model that uses incremental minimization to ensure the descent of the hydrogel free energy.

Using the stress-strain relationship, it can be demonstrated that Gent-type materials are strain-stiffening, while neo-Hookean materials exhibit strain-softening behavior. Moreover, the stress-strain relationship for hydrogels under plane stress condition with constant solvent concentration, barely differs from that of a purely elastic material. However, this changes in the approximation for plane strains. In addition, in case a solvent flow into or out of the hydrogel is allowed to describe the exchange with a surrounding solvent bath, then the stress-strain relationship and the stiffness of the material also differ significantly from that of a purely elastic material under plane stress condition.

Furthermore, turning towards cell migration and orientation on thin stretched hydrogel sheets, one initially recognizes expected behavior, i.e. cells orient themselves vertically, in the direction of stretch under small applied strain. However, when a critical value of the applied strain is reached, the orientation of the cells rapidly flips in the horizontal direction, which is an experimentally unexpected phenomenon, cf. [13]. The value of this critical strain depends on the strength of the traction force of the cells and is observed for both Gent-type and neo-Hookean hydrogels.

Moreover, at higher strains for non-conserved hydrogels under plane stress conditions, we observe a dramatic change in the morphology of the Gent hydrogel compared to the neo-Hookean one. While the Gent hydrogel undergoes a strong contraction and the solvent flows out of the hydrogel, the shape of the neo-Hookean hydrogel is only slightly contracted as a solvent flows into the material. This in turn has a significant effect on cell migration. In the case of the Gent hydrogel, the cells migrate towards the clamped boundaries and the formation of cell chains is suppressed. In the neo-Hookean model, on the other hand, the cells form horizontal long-range alignments that are distributed over the entire length of the stretched thin sheet.

In the current study we have not yet explored the range of traction forces imparted by a single cell where strain-stiffening effects become relevant. Thus, cells move rather passively and do not yet fully capture the mechanical reciprocity of the cell-hydrogel system. In order to investigate possible qualitative different behavior of the cell population for the Gent-type and neo-Hookean hydrogels in this regime, it will be necessary to allow for a higher spatial resolution near cells. This will also allow to robustly generate larger traction forces in the strain-stiffening range and thus cells become active agents. This will further entail modifications of the ABM that also accounts for hydrogel stiffness for example via a second derivative of the mechanical energy, for example as introduced in [57, 61, 62]. We note that our model framework for the cell-hydrogel system can be systematically extend the by adding, apart from further cell species, also further networks with prescribed properties, such as allowing for possible degradation, by adding a further phase into the formulation. Similarly, one can add another (non-conserved) phase to model the network fibers that are secreted by the cells and thus allow for remodeling of the hydrogel. In addition, further properties such as phase separation into regions of high and low fiber concentration induced by cell traction forces can be included. The systematic extensions of the model framework and their predictions form the basis for validation by *in vitro* experimental work in order to gain a fundamental understanding of the intricate mechanical interplay between cells and the ECM in tissue formation.

In general, the framework presented in this work provides a versatile toolbox to model complex processes that affect the cellular movement. Its true strength lies in extensibility, allowing for the integration of intricate biological processes such as biochemical signaling or cellular responses.

## Acknowledgment

AE and LS were supported by the German Research Foundation (DFG) under Germany’s Excellence Strategy – MATH^+^: The Berlin Mathematics Research Center (EXC-2046/1 – project ID: 390685689) via the project AA1-12^∗^. We acknowledge financial support by the DFG Priority Program SPP 2171 *Dynamic Wetting of Flexible, Adaptive, and Switchable Substrates* within the projects 422786086 (LS, AE) and 422792530 (DP). CD was supported by the DFG through the Collaborative Research Centre (CRC) 1444, project ID: 427826188.

## Footnotes

1 At this maximal value the elastic material self-intersects or 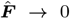 or 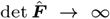, so that the solution of the mechanical problem becomes meaningless.

## References

[1] Janmey, P. A., Winer, J. P. & Weisel, J. W. Fibrin gels and their clinical and bioengineering applications. Journal of the Royal Society Interface 6, 1–10 (2009).

[2] Borgiani, E., Duda, G. N. & Checa, S. Multiscale modeling of bone healing: toward a systems biology approach. Frontiers in Physiology 8, 287 (2017).

[3] Hogrebe, N. J., Reinhardt, J. W. & Gooch, K. J. Biomaterial microarchitecture: a potent regulator of individual cell behavior and multicellular organization. Journal of Biomedical Materials Research Part A 105, 640–661 (2017).

[4] Brauer, E. et al. Collagen fibrils mechanically contribute to tissue contraction in an in vitro wound healing scenario. Advanced Science 6, 1801780 (2019).

[5] Checa, S., Rausch, M. K., Petersen, A., Kuhl, E. & Duda, G. N. The emergence of extracellular matrix mechanics and cell traction forces as important regulators of cellular selforganization. Biomechanics and Modeling in Mechanobiology 14, 1–13 (2015).

[6] Checa, S. & Prendergast, P. J. A mechanobiological model for tissue differentiation that includes angiogenesis: a lattice-based modeling approach. Annals of Biomedical Engineering 37, 129–145 (2009).

[7] Wang, J. H. & Grood, E. S. The strain magnitude and contact guidance determine orientation response of fibroblasts to cyclic substrate strains. Connective Tissue Research 41, 29–36 (2000).

[8] Isomursu, A. et al. Directed cell migration towards softer environments. Nature Materials 21, 1081–1090 (2022).

[9] Winer, J. P., Oake, S. & Janmey, P. A. Non-linear elasticity of extracellular matrices enables contractile cells to communicate local position and orientation. PloS one 4, e6382 (2009).

[10] Wen, Q. & Janmey, P. A. Effects of nonlinearity on cell–ECM interactions. Experimental Cell Research 319, 2481–2489 (2013).

[11] Grekas, G. et al. Cells exploit a phase transition to mechanically remodel the fibrous extracellular matrix. Journal of the Royal Society Interface 18, 20200823 (2021).

[12] Reinhardt, J. W., Krakauer, D. A. & Gooch, K. J. Complex matrix remodeling and durotaxis can emerge from simple rules for cellmatrix interaction in agent-based models. Journal of Biomechanical Engineering 135, 071003 (2013).

[13] Ahearne, M. Introduction to cell–hydrogel mechanosensing. Interface Focus 4, 20130038 (2014).

[14] Dazzi, C. et al. External mechanical loading overrules cell-cell mechanical communication in sprouting angiogenesis during early bone regeneration. PLOS Computational Biology 19, 1–27 (2023). URL 10.1371/journal.pcbi.1011647.

[15] Alisafaei, F., Chen, X., Leahy, T., Janmey, P. A. & Shenoy, V. B. Long-range mechanical signaling in biological systems. Soft matter 17, 241–253 (2021).

[16] Natan, S., Koren, Y., Shelah, O., Goren, S. & Lesman, A. Long-range mechanical coupling of cells in 3D fibrin gels. Molecular Biology of the Cell 31, 1474–1485 (2020).

[17] Liu, K., Wiendels, M., Yuan, H., Ruan, C. & Kouwer, P. H. Cell-matrix reciprocity in 3D culture models with nonlinear elasticity. Bioactive Materials 9, 316–331 (2022).

[18] Licup, A. J. et al. Stress controls the mechanics of collagen networks. Proceedings of the National Academy of Sciences 112, 9573–9578 (2015). URL 10.1073/pnas.1504258112.

[19] Han, Y. L. et al. Cell contraction induces long-ranged stress stiffening in the extracellular matrix. Proceedings of the National Academy of Sciences 115, 4075–4080 (2018).

[20] Haupert, S., Guérard, S., Mitton, D., Peyrin, F. & Laugier, P. Quantification of nonlinear elasticity for the evaluation of submillimeter crack length in cortical bone. Journal of the Mechanical Behavior of Biomedical Materials 48, 210–219 (2015).

[21] Reinhardt, J. W. & Gooch, K. J. An agent-based discrete collagen fiber network model of dynamic traction force-induced remodeling. Journal of Biomechanical Engineering 140 (2018).

[22] Wolf, K. et al. Collagen-based cell migration models in vitro and in vivo. Seminars in Cell & Developmental Biology 20, 931–941 (2009). URL https://www.sciencedirect.com/science/article/pii/S108495210900161X. Imaging in Cell and Developmental Biology Planar Cell Polarity.

[23] Caccavo, D., Cascone, S., Lamberti, G. & Barba, A. Hydrogels: experimental characterization and mathematical modelling of their mechanical and diffusive behaviour. Chemical Society Reviews 47, 2357–2373 (2018).

[24] Solbu, A. A. et al. Assessing cell migration in hydrogels: An overview of relevant materials and methods. Materials Today Bio 100537 (2022).

[25] Hazur, J., Endrizzi, N., Schubert, D. W., Boccaccini, A. R. & Fabry, B. Stress relaxation amplitude of hydrogels determines migration, proliferation, and morphology of cells in 3-D culture. Biomaterials Science 10, 270–280 (2022).

[26] Chaudhuri, O. Viscoelastic hydrogels for 3D cell culture. Biomaterials science 5, 1480–1490 (2017).

[27] Münster, S. et al. Strain history dependence of the nonlinear stress response of fibrin and collagen networks. Proceedings of the National Academy of Sciences 110, 12197–12202 (2013). URL 10.1073/pnas.1222787110.

[28] Abdalrahman, T. & Checa, S. On the role of mechanical signals on sprouting angiogenesis through computer modeling approaches. Biomechanics and Modeling in Mechanobiology 21, 1623–1640 (2022). URL 10.1007/s10237-022-01648-4.

[29] Borgiani, E., Duda, G. N., Willie, B. M. & Checa, S. Bone morphogenetic protein 2-induced cellular chemotaxis drives tissue patterning during critical-sized bone defect healing: an in silico study. Biomechanics and Modeling in Mechanobiology 20, 1627–1644 (2021). URL 10.1007/s10237-021-01466-0.

[30] Perier-Metz, C., Duda, G. N. & Checa, S. Initial mechanical conditions within an optimized bone scaffold do not ensure bone regeneration –an in silico analysis. Biomechanics and Modeling in Mechanobiology 20, 1723–1731 (2021). URL 10.1007/s10237-021-01472-2.

[31] Balzani, D., Heinlein, A., Klawonn, A. & et al. Comparison of arterial wall models in fluid–structure interaction simulations. Comput Mech 72, 949–965 (2023).

[32] Peirlinck, M., Linka, K., Hurtado, J. A. & Kuhl, E. On automated model discovery and a universal material subroutine for hyperelastic materials. Computer Methods in Applied Mechanics and Engineering 418, 116534 (2024). URL https://www.sciencedirect.com/science/article/pii/S0045782523006588.

[33] Ricker, A., Gierig, M. & Wriggers, P. Multiplicative, Non-Newtonian Viscoelasticity Models for Rubber Materials and Brain Tissues: Numerical Treatment and Comparative Studies. Arch Computat Methods Eng 30, 2889–2927 (2023).

[34] Hennessy, M. G., Münch, A. & Wagner, B. Phase separation in swelling and deswelling hydrogels with a free boundary. Physical Review E 101, 032501 (2020).

[35] Schmeller, L. & Peschka, D. Gradient Flows for Coupling Order Parameters and Mechanics. SIAM Journal on Applied Mathematics 83, 225–253 (2023).

[36] Huggins, M. L. Theory of solutions of high polymers. Journal of the American Chemical Society 64, 1712–1719 (1942).

[37] Flory, P. J. Thermodynamics of high polymer solutions. The Journal of chemical physics 10, 51–61 (1942).

[38] Bosnjak, N., Nadimpalli, S., Okumura, D. & Chester, S. A. Experiments and modeling of the viscoelastic behavior of polymeric gels. Journal of the Mechanics and Physics of Solids 137, 103829 (2020).

[39] Rognes, M. E., Calderer, M.-C. & Micek, C. A. Modelling of and mixed finite element methods for gels in biomedical applications. SIAM Journal on Applied Mathematics 70, 1305–1329 (2009).

[40] Kim, B. et al. A comparison among Neo-Hookean model, Mooney-Rivlin model, and Ogden model for chloroprene rubber. International Journal of Precision Engineering and Manufacturing 13, 759–764 (2012).

[41] Gent, A. N. A new constitutive relation for rubber. Rubber Chemistry and Technology 69, 59–61 (1996).

[42] Jin, L. & Suo, Z. Smoothening creases on surfaces of strain-stiffening materials. Journal of the Mechanics and Physics of Solids 74, 68–79 (2015).

[43] Holzapfel, G. Biomechanics of soft tissue, 1049–1063 (Academic Press, United States, 2001), volume iii, multiphysics behaviors, chapter 10, composite media edn.

[44] Dunn, M. G., Silver, F. H. & Swann, D. A. Mechanical Analysis of Hypertrophic Scar Tissue: Structural Basis for Apparent Increased Rigidity. Journal of Investigative Dermatology 84, 9–13 (1985).

[45] Imaoka, C. et al. Inverse mechanical-swelling coupling of a highly deformed double-network gel. Science Advances 9, eabp8351 (2023).

[46] Alnæs, M. et al. The FEniCS project version 1.5. Archive of Numerical Software 3 (2015).

[47] Logg, A., Mardal, K.-A. & Wells, G. Automated solution of differential equations by the finite element method: The FEniCS book Vol. 84 (Springer Science & Business Media, 2012).

[48] Erhardt, A. H., Peschka, D. & Schmeller, L. Modeling cellular self-organization in strain-stiffening hydrogels. https://github.com/andreerhardt/hydrogel abm (2023).

[49] Peng, Q. Mathematical Aspects of Cell-Based and Agent-Based Modelling for Skin Contraction after Deep Tissue Injury. Ph.D. thesis, Delft University of Technology (2021).

[50] Westman, A. M., Peirce, S. M., Christ, G. J. & Blemker, S. S. Agent-based model provides insight into the mechanisms behind failed regeneration following volumetric muscle loss injury. PLOS Computational Biology 17, 1–29 (2021). URL 10.1371/journal.pcbi.1008937.

[51] Wong, S., Guo, W.-H. & Wang, Y.-L. Fibroblasts probe substrate rigidity with filopodia extensions before occupying an area. Proceedings of the National Academy of Sciences 111, 17176–17181 (2014).

[52] Cerda, E. & Mahadevan, L. Geometry and physics of wrinkling. Physical Review Letters 90, 074302 (2003).

[53] Wang, T., Yang, Y. & Xu, F. Mechanics of tension-induced film wrinkling and restabilization: a review. Proceedings of the Royal Society A 478, 20220149 (2022).

[54] Ideses, Y. et al. Spontaneous buckling of contractile poroelastic actomyosin sheets. Nature Communications 9, 2461 (2018).

[55] Zakharov, A. & Dasbiswas, K. Modeling mechanochemical pattern formation in elastic sheets of biological matter. The European Physical Journal E 44, 82 (2021).

[56] Livne, G., Gat, S., Armon, S. & Bernheim-Groswasser, A. Artificial contractile actomyosin gels recreate the curved and wrinkling shapes of cells and tissues. bioRxiv 2023–03 (2023).

[57] Steinwachs, J. et al. Three-dimensional force microscopy of cells in biopolymer networks. Nature Methods 13, 171–176 (2016).

[58] Van Helvert, S. & Friedl, P. Strain stiffening of fibrillar collagen during individual and collective cell migration identified by AFM nanoindentation. ACS Applied Materials & Interfaces 8, 21946–21955 (2016).

[59] Jaspers, M. et al. Ultra-responsive soft matter from strain-stiffening hydrogels. Nature Communications 5, 1–8 (2014).

[60] Jaspers, M. et al. Nonlinear mechanics of hybrid polymer networks that mimic the complex mechanical environment of cells. Nature communications 8, 1–10 (2017).

[61] Steinwachs, J. Cellular Forces during Migration through Collagen Networks (Friedrich-Alexander-Universitaet Erlangen-Nuernberg (Germany), 2015).

[62] Cóndor, M. et al. Breast cancer cells adapt contractile forces to overcome steric hindrance. Biophysical Journal 116, 1305–1312 (2019).

